# NMR analysis of the homodimeric structure of a D-Ala-D-Ala metallopeptidase, VanX, from vancomycin-resistant bacteria

**DOI:** 10.1101/2023.10.13.562161

**Authors:** Tsuyoshi Konuma, Tomoyo Takai, Chieko Tsuchiya, Masayuki Nishida, Miyu Hashiba, Yudai Yamada, Ayaka Oe, Haruka Shirai, Aritaka Nagadoi, Eriko Chikaishi, Ken-ichi Akagi, Satoko Akashi, Toshio Yamazaki, Hideo Akutsu, Takahisa Ikegami

## Abstract

Bacteria that have acquired resistance to most antibiotics, particularly those causing nosocomial infections, create serious problems. Among these, the emergence of vancomycin-resistant enterococci was a tremendous shock considering that vancomycin is the last resort for controlling methicillin-resistant *Staphylococcus aureus*. Therefore, there is an urgent need to develop an inhibitor of VanX, a protein involved in vancomycin resistance. Although the crystal structure of VanX has been resolved, its asymmetric unit contains six molecules aligned in a row. We have developed a structural model of VanX as a stable dimer in solution, primarily utilizing nuclear magnetic resonance (NMR) residual dipolar coupling. Despite the 46 kDa molecular mass of the dimer, which is typically considered too large for NMR studies, we successfully assigned the main chain using an amino acid-selective unlabeling method. Because we found that the Zn^2+^-coordinating active sites in the dimer structure were situated in the opposite direction to the dimer interface, we generated an active monomer by mutating the dimer interface. The monomer consists of only 202 amino acids and is expected to be used in future studies to screen and improve inhibitors using NMR.

## Introduction

Nosocomial bacterial infections resistant to many types of antibiotics have become a major social problem. A publication described that the mortality associated with bacterial antimicrobial resistance amounted to nearly 4.95 million cases in 2019, with projections indicating an escalation to 10 million cases annually by the year 2050 ^1^. Vancomycin, a glycopeptide antibiotic, was developed in 1956 to combat these bacteria. This drug was once considered a last resort because it affects methicillin-resistant *Staphylococcus aureus* (MRSA), which is resistant to many other known antibiotics. However, in 1988, vancomycin-resistant *Enterococci faecium* (VREF) emerged ^2^. Postoperatively, VREF poses a serious threat to patients with low or suppressed immunity.

Bacterial cell walls are constructed of peptidoglycan, which is formed by the cross-linking of pentapeptides, L-Ala^1^-D-Glu^2^-L-Lys^3^-D-Ala^4^-D-Ala^5^, bound to N-acetylmuramic acid (for *enterococci*) ^3^. Vancomycin interferes with bacterial cell wall synthesis by binding to the D-Ala^4^-D-Ala^5^ moiety before cross-linking occurs, causing the lysis of gram-positive bacteria. VanX (UniProt: A0A286KC57_ENTAV) functions as a D-alanyl-D-alanine (D-Ala-D-Ala) dipeptidase, and another enzyme, VanA, ligates D-lactate (D-Lac) to the hydrolyzed product to yield a depsipeptide, D-Ala^4^-D-Lac^5^ ^4,5^. Since vancomycin has a thousand-fold reduced affinity for the D-Ala^4^-D-Lac^5^ moiety, it no longer inhibits the synthesis of peptidoglycan cell walls. In contrast, D-Ala-D-Lac does not inhibit the peptidoglycan synthesis.

The structure of VanX derived from *E. faecium* was determined by X-ray crystallography ^6^. The active site of VanX coordinates the zinc ion (Zn^2+^) ^7^ through His116, Asp123, Glu181, and His184. Glu181 coordinates with Zn^2+^ via a hydrogen bond with a water molecule. Upon recognizing D-Ala-D-Ala, Glu181 acts as a catalytic base that activates water by extracting a proton from it, and VanX hydrolyzes the substrate into two D-Ala monomers ^8–11^.

Several vanX inhibitors have been developed to suppress vancomycin resistance. Inhibitory activities have been shown by a few phosphonate ^12^, phosphinate ^13–15^, and phosphonamidate ^16,17^ analogs of tetrahedral adducts that mimic the transition state for the hydrolysis of D-Ala-D-Ala. However, none of these inhibitors can pass through the membrane to reach the bacterial cytoplasm and thus have not been considered effective inhibitors. Recently, Muthyala *et al*. developed two cyclic thiohydroxamic acid-based peptide analogs that could penetrate cells and exhibited low-micromolar inhibitory activity against VREF in a cell-based assay ^18^.

The crystal structure of VanX was solved; however, according to the registered coordinates (PDB:1r44), the asymmetric unit consists of six molecules aligned in a row ^6^. Herein, we propose a model structure for VanX that exists stably in solution as a dimer. The dimer structure was determined by nuclear magnetic resonance (NMR) residual dipole coupling (RDC) and transfer cross-saturation (TCS). VanX exhibited a homodimeric structure with a molecular mass of 46,760 Da, where each subunit contributed a mass of 23,380 Da (202 amino acids for each), which made it difficult to analyze by NMR; however, mutation of the two cysteine residues to serine (C78/157S) and deuteration made it possible to assign the main-chain nuclei. We succeeded in splitting the subunits using an additional mutation (C78/157S, W24R) and preparing the monomer. As a result, the molecular weight was halved, making NMR analysis more accessible. Our basic data will be useful for future development of VanX inhibitors using NMR.

## Results

### Sample preparation

The expression of soluble VanX in *Escherichia coli* (*E. coli*) bacteria led to host lysis 参考文献.

To prevent this, we strictly controlled the onset of expression by using the BL21 (DE3) pLysS strain. We also maintained the culture temperature at 37°C to prevent lysis. This is because VanX is often expressed as soluble in cultures at 15°C, but mostly forms inclusion bodies at 37°C ^19^. VanX was purified by refolding inclusion bodies. However, wild-type VanX aggregated and precipitated during NMR measurements despite attempting various solvent conditions, making it impossible to obtain a series of three-dimensional (3D) spectra for main-chain assignment. To overcome this problem, we mutated two cysteine residues (Cys78 and Cys157) to serine. This mutant was sufficiently stable to undergo NMR measurements at pH 4 and 7 (theoretical pI = 5.58). As previously reported ^20^, these two cysteine residues are not essential for the activity or zinc coordination of VanX. The purified VanX^C78/157S^ was subjected to analytical gel filtration. The results showed that VanX^C78/157S^ was eluted immediately after elution with a molecular weight marker (43 kDa) (Figure 1). This indicates that VanX^C78/157S^ exists stably as a dimer in solution (twice as much as 23 kDa). We also confirmed that the mutant exhibits the desired activity (described later). However, even without the cysteine residues, the mutant was still highly unstable at subunit concentrations greater than 0.1 mM. To obtain spectra with sufficient sensitivity, we purified four independent samples of VanX^C78/157S^ labeled with ^2^H, ^15^N, and ^13^C from each 1 L of culture medium. The spectra of the four samples were superimposed to achieve high signal-to-noise ratios.

**Figure 1.**
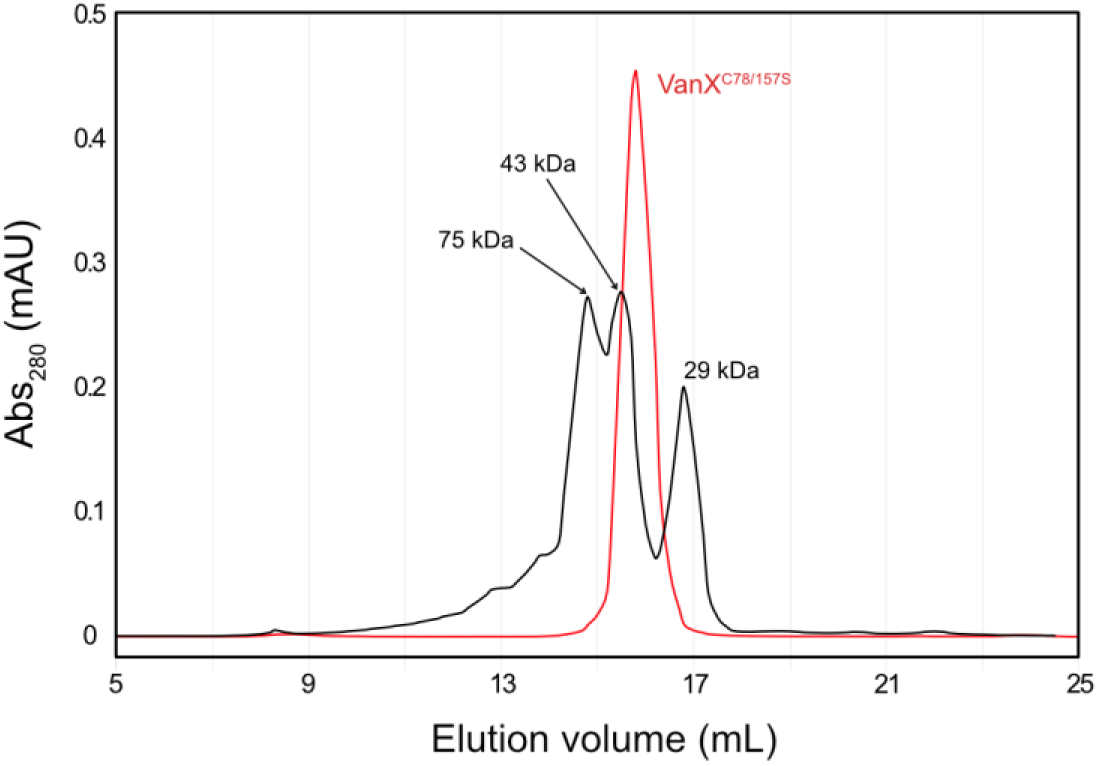
Molecular weight analysis of VanX^C78/157S^ by gel-filtration chromatography using a Superdex™ 200 10/300 GL column. A 20 mM Tris-HCl (pH 8.0) buffer containing 30 µM ZnCl_2_ and 150 mM NaCl was flowed at a rate of 0.5 mL/min. The elution profile of molecular weight markers (conalbumin: 75 kDa, ovalbumin: 43 kDa, and carbonic anhydrase: 29 kDa) is shown in black, while the elution profile of VanX^C78/157S^ is shown in red.

### NMR assignment using amino acid-selective unlabeling

A series of 3D TROSY-type spectra were recorded using [uniformly-^2^H, ^15^N, ^13^C]-VanX^C78/157S^ with a sensitive cryogenic probe. However, only 75% of the main-chains were assigned. To enhance this assignment, amino acid-selective unlabeling was employed. Specifically, we prepared [^15^N, ^13^C]-VanX^C78/157S^ in which each of the amino acids Lys, Arg, Asn, Ala, His, Gln, Met, Thr, Ser, and Tyr was unlabeled. We measured 2D ^1^H-^15^N TROSY-HSQC spectra at pH 4 and 7, with the spectra at pH 4 exhibiting higher spectral quality owing to the slower exchange of the amide ^1^H spins with the solvent water. To establish a connection between the assignment at pH 4 and 7, we measured 3D TROSY-HNCO spectra of 70 μM [^15^N, ^13^C]-VanX^C78/157S^ at pH 4, 5, 6, and 7 and followed the chemical shift changes with pH. This method facilitated the assignment of the respective amino acids as follows, with the expected numbers shown as denominators: Lys 9/9, Arg 14/14, Asn 10/11, Ala 14/15, His 5/5, Gln 4/4, Met 2/6, Thr 9/9, Ser 17/17, Trp 6/6, Gly 14/14, Tyr 13/13, and Phe 3/9 (Figures 2 and S1). It is well established that Thr undergoes metabolic scrambling with Gly as does Ser with Gly and Cys ^21–23^. Therefore, we assigned these residues by combining the spectra of the samples in which each of Thr and Ser was unlabeled (our VanX mutant contained no Cys residues). Additionally, the Trp peaks disappeared in the spectrum of the Ser-unlabeled sample. This is because Trp biosynthesis utilizes Ser as the main chain material after indole ring formation. Although 1.0 g/L Tyr was added to the medium, unlabeled Tyr did not eliminate the corresponding peaks. This is likely due to the low solubility of Tyr (only 0.45 g/L). Finally, we assigned 95.0 and 97.5% of the backbone amide ^1^H/^15^N signals at pH 7 and 4, respectively (Figures 3 and S2).

**Figure 2.**
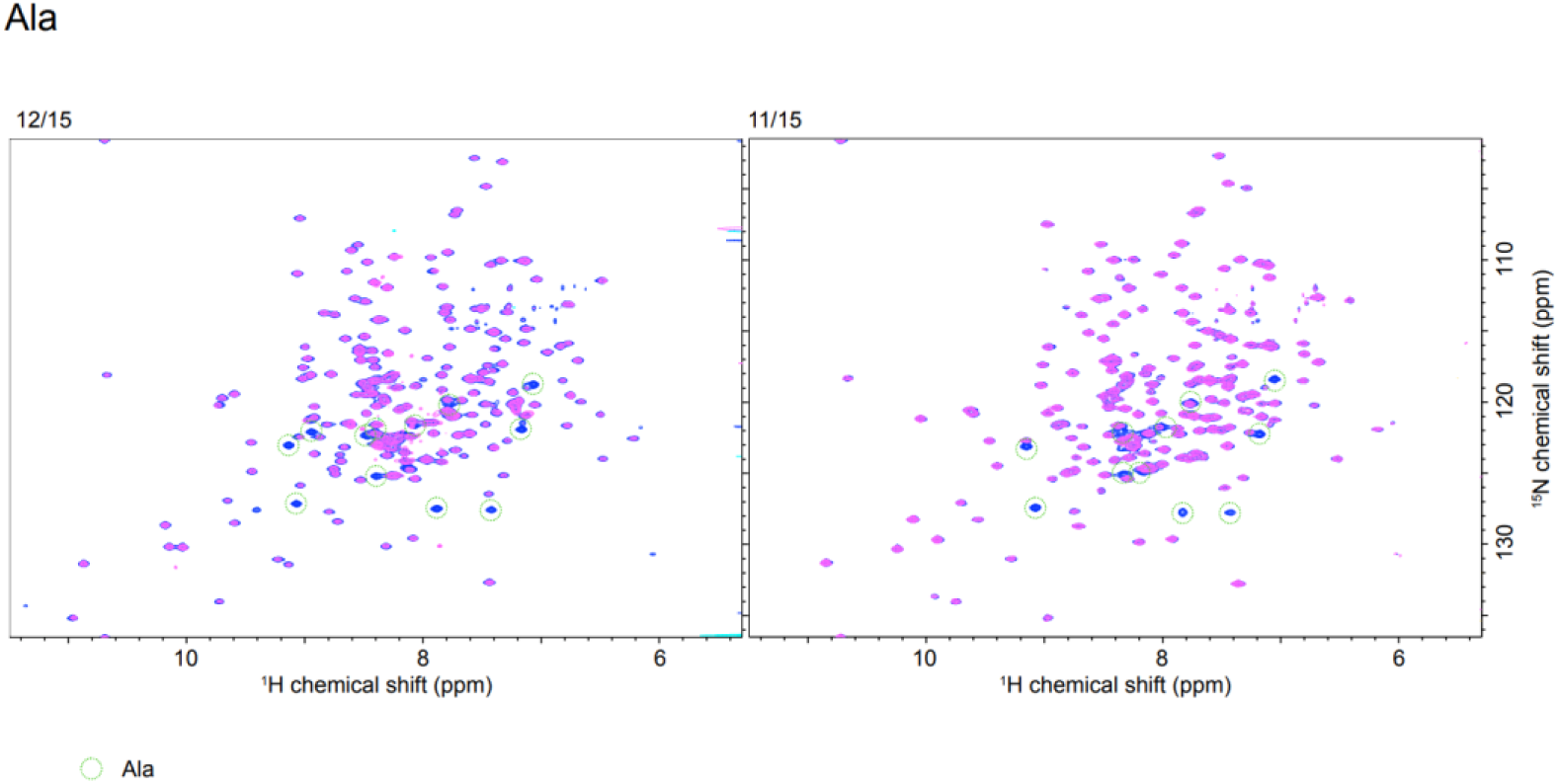
Overlay of 2D ^1^H-^15^N TROSY-HSQC spectra of reference VanX and VanX with alanines unlabeled. The pH 4 and pH 7 spectra are on the left and right sides, respectively. Green circles indicate unlabeled target amino acids. The fraction n/m, written on the left side of each spectrum, indicates that n out of the expected m peaks were actually broadened. Similar figures for the other specific amino acid unlabeling can be referenced in Figure S1.

**Figure 3.**
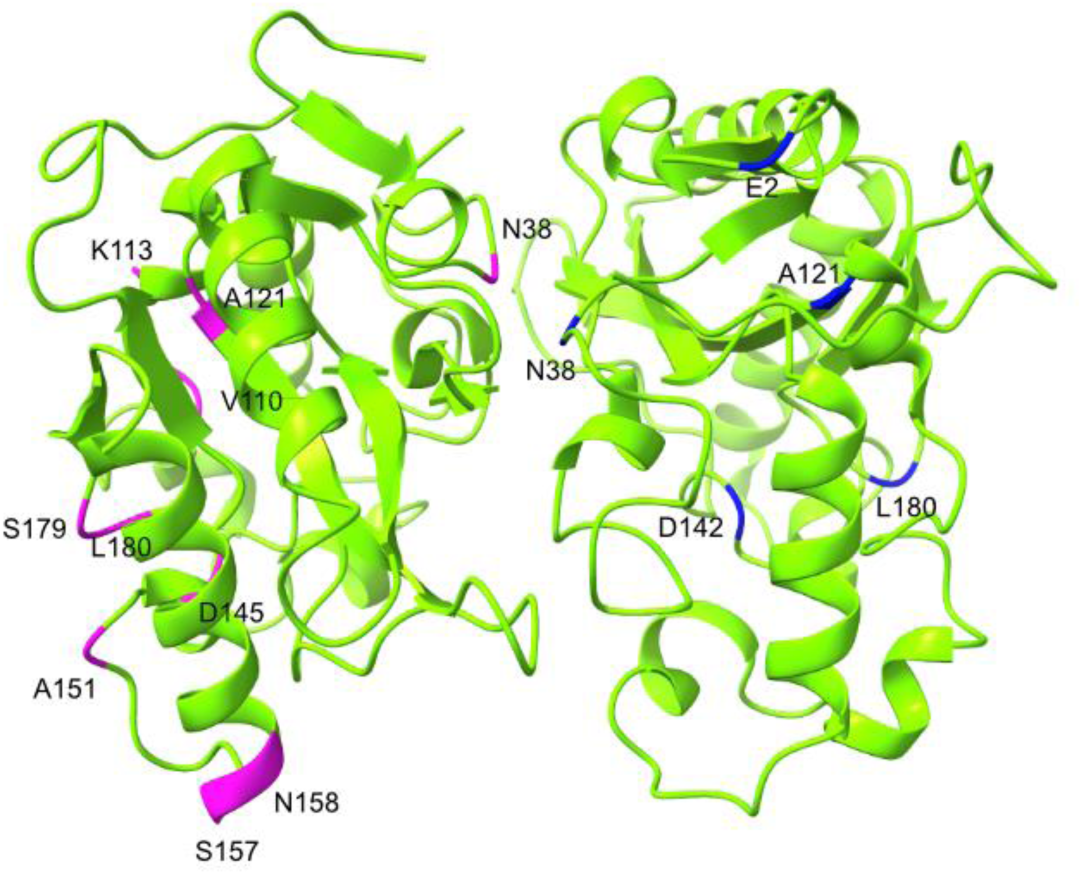
Location of unassigned residues in the 3D structure. The residues whose amide ^1^H/^15^N peaks were not assigned at pH 7 and 4 are depicted in magenta and blue on the B (left) and C (right) subunits of the crystal structure (1r44), respectively. Diagrams of the protein structures were created using ChimeraX ^24^.

### Detection of subunit interaction by TCS

In the transfer cross saturation (TCS) experiments, continuous pulses were applied to the methyl groups of a protonated [^13^C]-VanX subunit to saturate the ^1^H spin polarization of the entire subunit. Because the side chains of the other [^15^N, ^2^H]-labeled subunit were deuterated, only the [^13^C]-labeled subunit was directly affected by the pulses. Saturation was then transferred to the ^1^H^N^ amide groups at the subunit interface of the [^15^N, ^2^H]-labeled subunit. The interaction region was identified based on the decreased signal intensity of the saturated ^1^H^N^ spins. The ^1^H^N^ signals were subsequently edited using the ^1^H-^15^N-HSQC sequence, which allowed the detection of only the saturation transferred from the [^13^C]-labeled subunit to the [^15^N, ^2^H]-labeled subunit in a homodimer. Figure 4 shows the amino acids that exhibited a marked decrease in peak intensity. These amino acids are R16, W17, D18, A19, R39, L65, and L66 (red). The other amino acids that exhibited minor decreases in peak intensity were W24, G34, V37, R128, L129, L134, and V201 (pink). Mapping of these residues onto the X-ray crystal structure of VanX (PDB:1r44) revealed that the subunit B/C model, which shares the same dimer conformation as subunit D/E, aligned with the results obtained from our TCS experiment.

**Figure 4.**
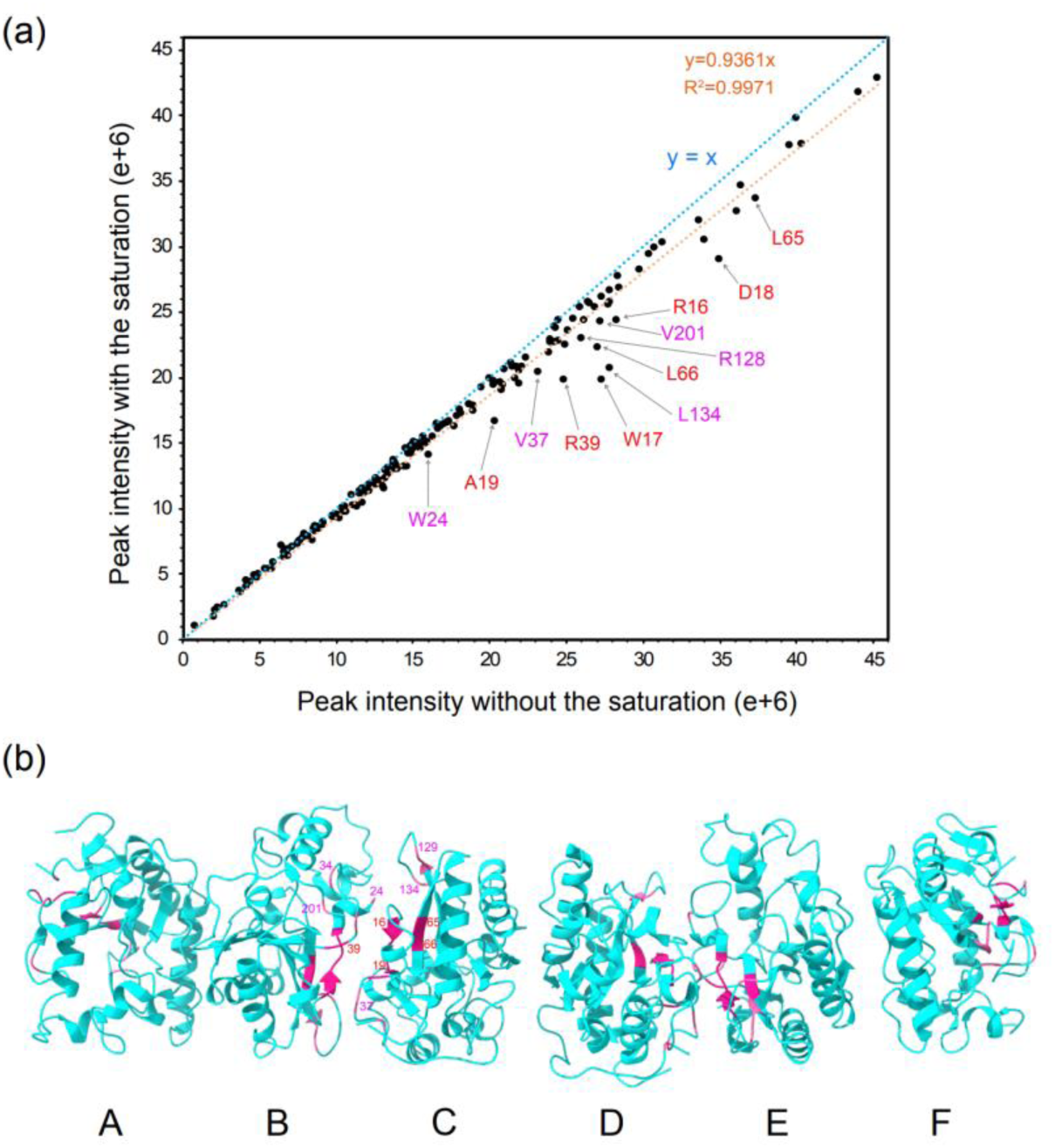
Results of the transfer cross saturation (TCS) experiments. (a) The graph shows the peak intensity of ^1^H^N^ in the TCS experiments. Four different combinations of saturation pulse intensity (100 and 200 Hz) and duration (0.5, 1.0, and 2.0 s) were used. The graph represents the data obtained with a saturation pulse intensity of 100 Hz and a duration of 2.0 s, which was consistent with the overall findings. (b) Amino acids that showed a decrease in peak intensity less than 85% under all four conditions are highlighted in red on the crystal structure. Additionally, amino acids that underwent a decrease in peak intensity under two or three of the four conditions are indicated in pink. On the crystal structure of VanX (1r44), the six monomers in the asymmetric unit are labeled as A through F.

### Detection of inter-subunit residues by NOE

The combination of [^13^C]-VanX and [^2^H, ^15^N]-VanX is applicable not only for TCS but also for inter-subunit nuclear *Overhauser* effect (NOE) experiments. We attempted 4D ^13^C-HSQC-NOESY-^15^N-HSQC measurements; however, low sensitivity prevented signal detection. Next, we detected NOEs, including the backbone ^1^H^N^, using [^13^C, ^15^N, partially ^2^H]-VanX, with a particular focus on the side chains of Trp, because these residues were found in the subunit interface of model B/C. The protein was expressed in *E. coli* cultured in D_2_O-based M9 medium supplemented with ^15^NH_4_Cl and [^13^C]-glucose. The incorporation of deuterons from D_2_O during protein biosynthesis results in the partial substitution of non-labile protons derived from glucose. Although this substitution leads to a decrease in proton density within the proteins, thereby reducing their absolute sensitivity, the benefits outweigh this drawback as it enhances the intensity of the NOE.

Bidirectional NOEs were detected between the side-chain amide group of a specific Trp residue and the main-chain amide group of L134 (Figure 5). Subsequently, in the 2D ^1^H-^15^N heteronuclear single quantum coherence (HSQC) spectrum of the W24R mutant (Figure 11), the peak corresponding to the Trp side chain disappeared, confirming that it originated from Trp24. This finding suggests spatial proximity between Trp24 and Leu134. Because these two residues are located distantly within a subunit, the observed NOE automatically indicates an intersubunit interaction. Notably, Bussiere *et al*., who solved the crystal structure of VanX, noticed that Trp24 protruded from the subunit surface, and they expected that this would contribute to inter-subunit interactions ^6^. Although the TCS results established an inter-subunit contact interface, they were unable to determine their precise relative orientation. In this context, NOE considerably restricted the relative orientation, consequently increasing the likelihood of the B/C model.

**Figure 5.**
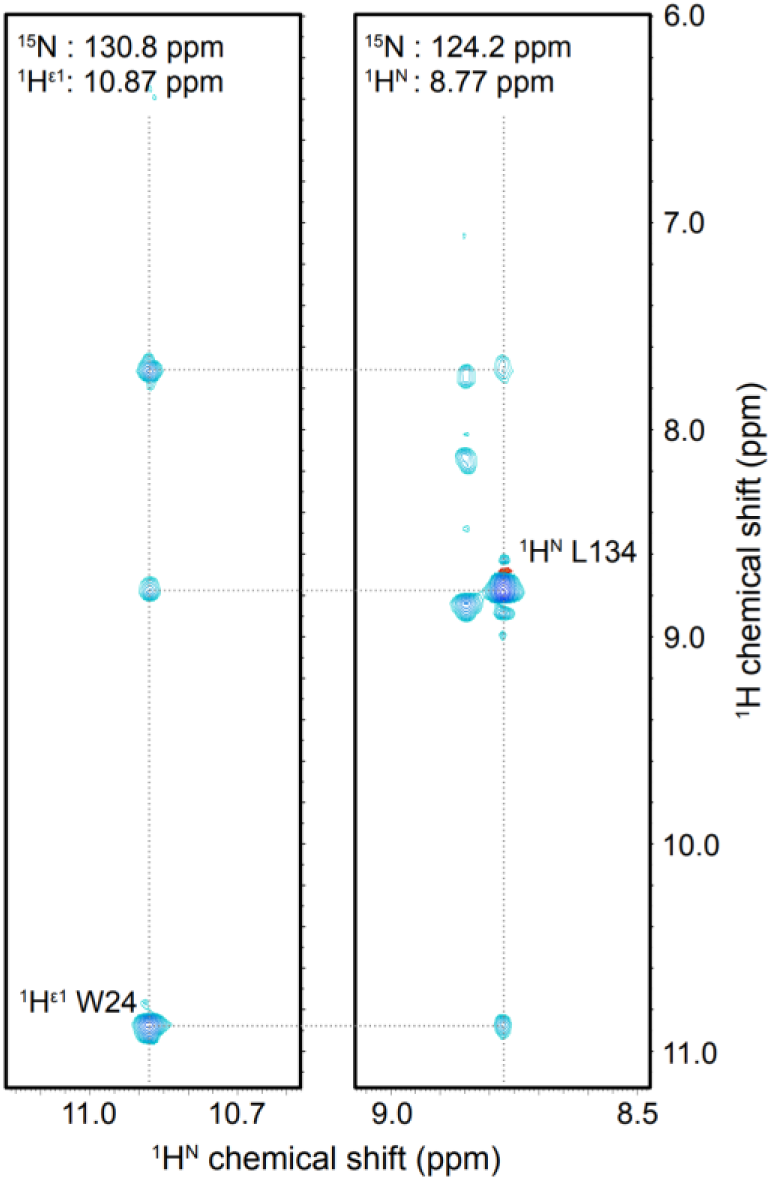
Strips (^1^H^N^/^1^H) of a 3D NOESY-^15^N-TROSY-HSQC spectrum. Results were acquired using 0.19 mM [^15^N, ^13^C, partially ^2^H]-VanX^C78/157S^ in a 50 mM KCl buffer (pH 7.0) supplemented with 30 μM ZnCl_2_ and 10% D_2_O, at a ^1^H resonance frequency of 800 MHz and a temperature of 303 K, using a mixing time of 120 ms. The left and right strips exhibited NOE peaks corresponding to the side chain ^1^H^ε1^ of Trp24 and the main chain ^1^H^N^ of Leu134, respectively. The diagonal peaks were assigned accordingly. The ^1^H direct (horizontal) measurement axis is for TROSY peaks. The chemical shift values on the ^1^H indirect (vertical) measurement axis were adjusted by −^1^*J*/2 (= 46.5 Hz) to match both dimensions.

### Determination of the relative orientation between subunits by RDC

The TCS method facilitated the identification of specific amino acid residues located at the dimer interface. Moreover, the NOE analysis narrowed down the potential homodimeric structure. However, the structural constraints derived from this information alone are insufficient to determine the exact orientation between the two subunits. For this purpose, information derived from residual dipolar couplings (RDCs) was incorporated. Initially, we measured the 2D ^1^H-^15^N TROSY and anti-TROSY spectra of [^2^H, ^15^N, ^13^C]-VanX^C78/157S^, which was aligned with the static magnetic field *B*_0_, using the coat protein of the *Pf1* filamentous bacteriophage. By combining these spectra, we obtained the difference in ^15^N resonance values for each residue (^1^*J*_HN_+^1^*D*_HN_), where ^1^*J*_HN_ and ^1^*D*_HN_ denote the scalar and RDC constants of the amide ^1^H/^15^N, respectively. (Figure 6). In addition, we extracted counterparts from an unoriented isotropic sample (^1^*J*_HN_). Finally, we calculated the RDC for each residue (^1^*D*_HN_) by obtaining the difference between the two split widths. The RDC values observed for magnetically oriented proteins depend on the angle formed by the corresponding amide ^1^H/^15^N bond relative to the static magnetic field *B*_0_ (Figure S3).

**Figure 6.**
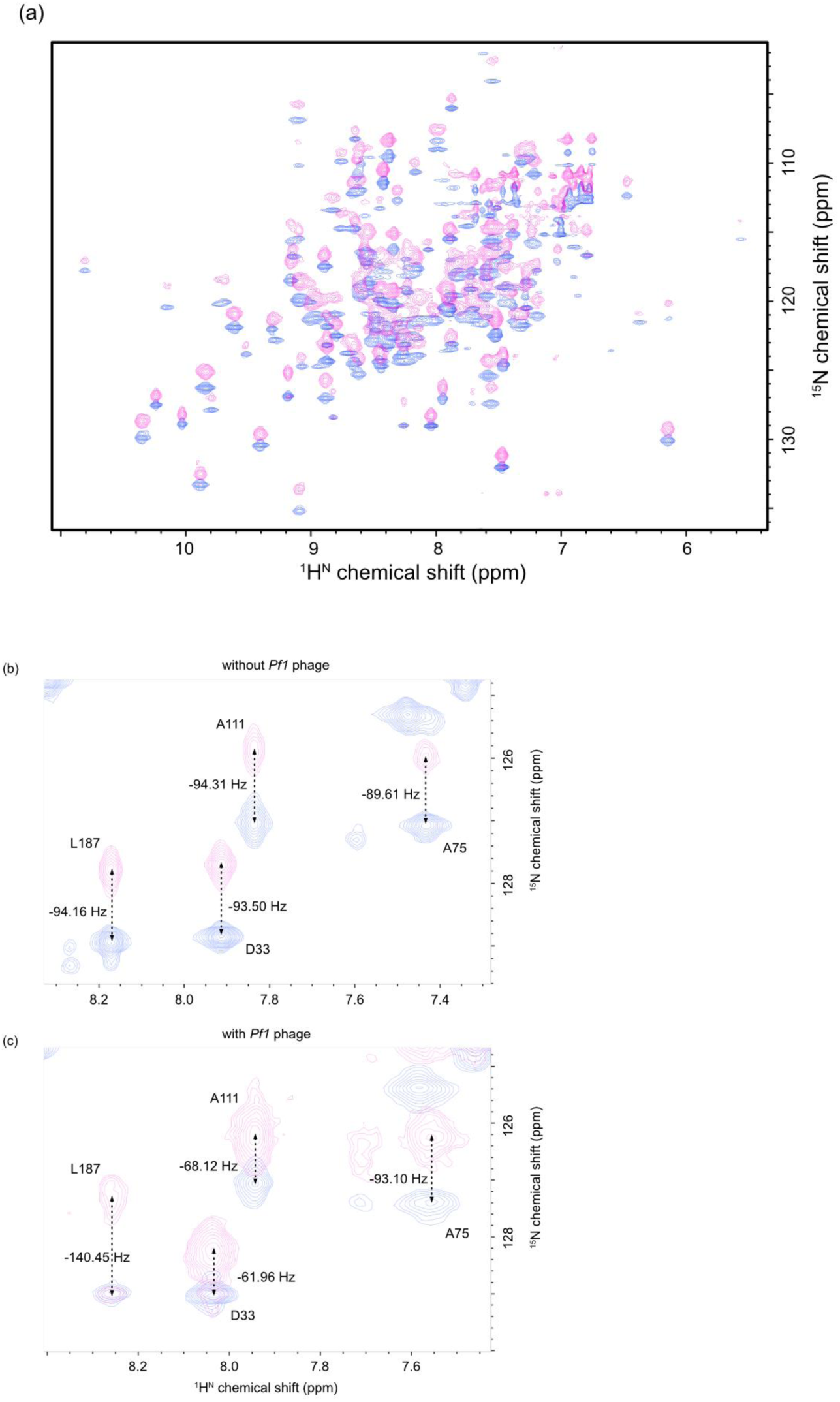
Overlay of the 2D ^1^H-^15^N TROSY (blue) and anti-TROSY (magenta) spectra obtained from [^2^H, ^15^N, ^13^C]-VanX^C78/157S^, which was aligned along the static magnetic field using the *Pf1*-phage coat proteins (a). Enlarged view of the superposition of spectra in the absence (b) and presence (c) of *Pf1*-phage coat proteins. The four assigned ^1^H/^15^N amide peaks in the figure are split along the ^15^N dimension, as indicated by the arrows and values. The chemical shift values are not adjusted between the spectra (b) and (c). Their displacement is attributed to the different D_2_O lock signals in the presence (c) and absence (b) of ^2^H residual quadrupolar coupling.

### Calculation of the homodimer structure

The dimer conformations were calculated using HADDOCK 2.4 on the basis of the “Modelling a homo-oligomeric complex from MS cross-links” procedure described in a *bonvin-lab* webpage (https://www.bonvinlab.org/education/HADDOCK24/XL-MS-oligomer/). The initial alignment tensor values (*D*a and *D*r) were estimated to be −1.27e-03 and −1.75e-04, respectively, using the Pales software (Figure 7) 25. These values were then scaled by a factor of 21,700 and supplied as input to HADDOCK (*i.e.*, −27.5 Hz (*D*=*D*a) and 0.136 (*R*=*D*r/*D*a), respectively). In total, 132 RDC-based orientational restraints and seven TCS-based intersubunit ambiguous restraints were imposed in the calculations. The latter restraints involved residues in segments B (selected from residues 16, 17, 18, 19, 39, 65, and 66, exhibiting remarkable TCS effects) and C (selected from residues 16, 17, 18, 19, 39, 65, 66, 24, 34, 37, 128, 129, 134, and 201, exhibiting both remarkable and minor TCS effects). These constraints were applied *c*_2_-symmetrically between the subunits B and C in the calculations.

**Figure 7.**
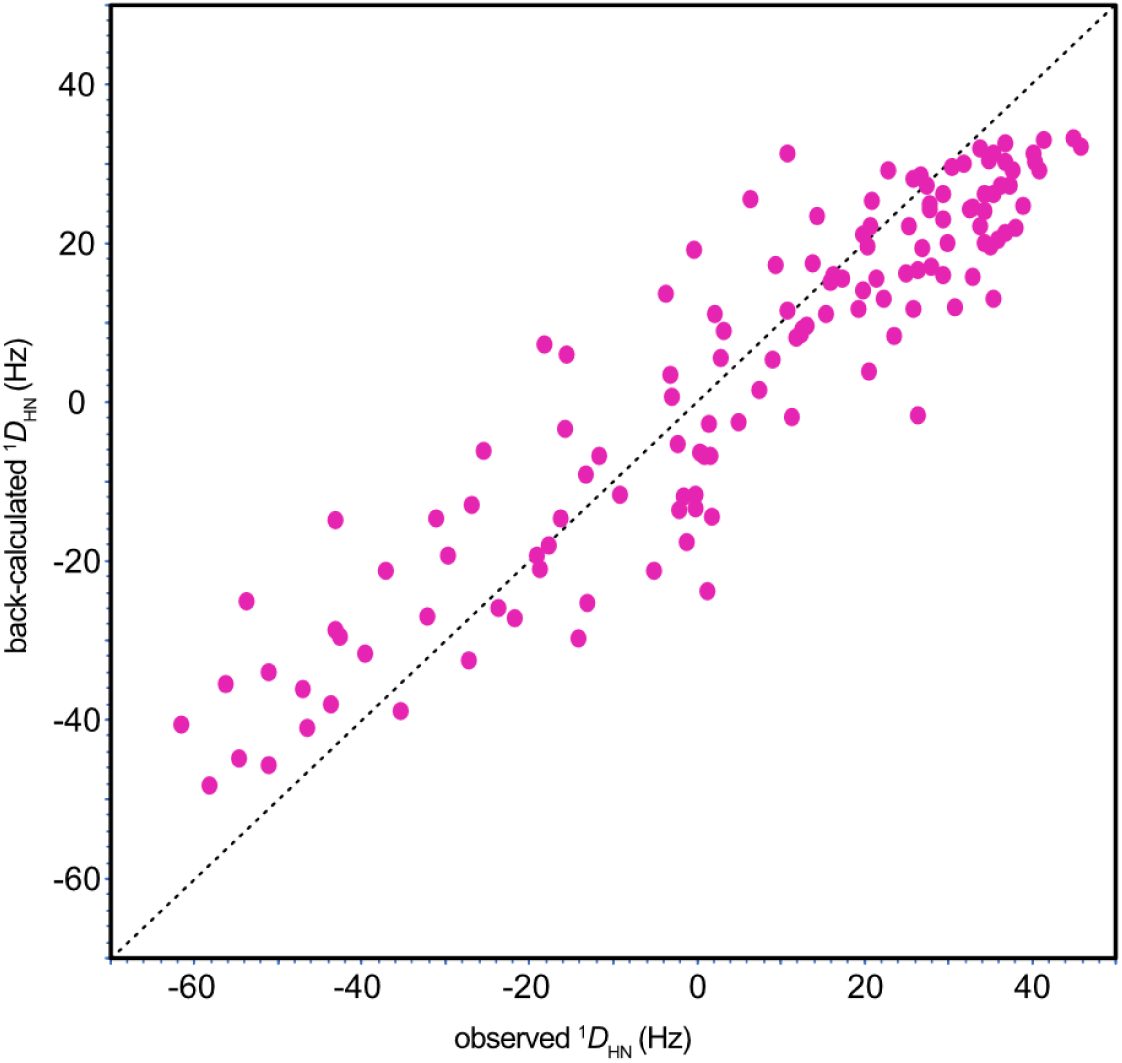
Correlation between the observed and back-calculated RDCs. The Pearson’s correlation coefficient was 0.88.

HADDOCK computations yielded 200 structures, all of which were classified into a single cluster (Table 1). This finding confirmed that the structures of the cluster accurately reflected the experimental conditions. To further validate our results, we conducted a comparative analysis between the best structure and the B/C subunits’ structure in the registered crystal structure (1r44). When the B subunits were superimposed (aligned), the heavy atoms of the B and C subunits in the main-chain were displaced by 0.34 and 1.10 Å, respectively (Figures 8, S3, and S4). In conclusion, our computational approach yielded a structure that closely resembled the dimeric structure of the B/C subunits in the crystal structure.

**Figure 8.**
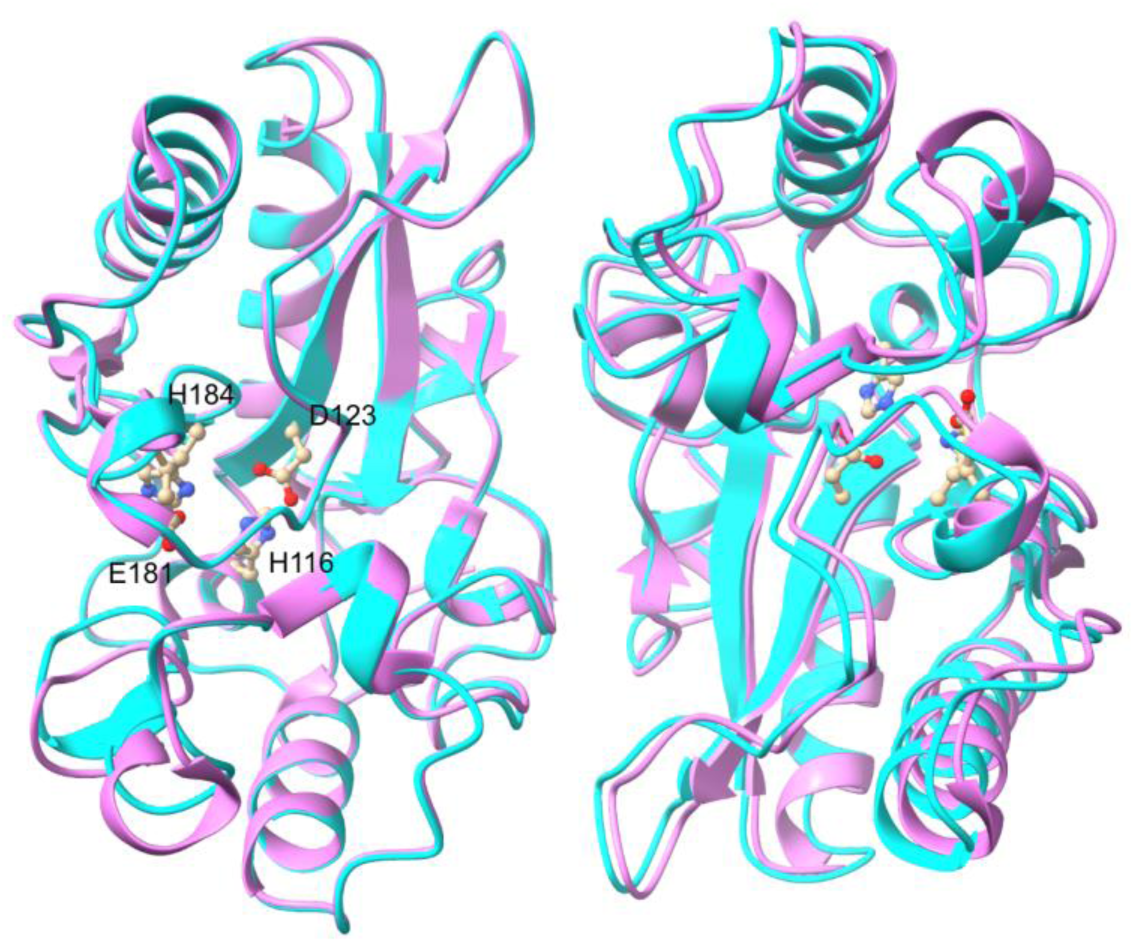
Comparison between the best structure, determined by HADDOCK (cyan), and the B/C subunits in the crystal structure (1r44) (pink). The N, C^α^, and C’ atoms in the backbones of the B subunits (left side) were superimposed. The side-chains around the active site and those in the subunit interface are illustrated using a ball-and-stick model.

**Table 1.** Results of structural calculations using HADDOCK 2.4. The output energy values are shown.

|  | The best cluster |
| --- | --- |
| HADDOCK score | 135.2 ± 4.8 |
| Cluster size (/200) | 200 |
| RMSD* from the overall lowest-energy structure | 0.4 ± 0.3 |
| Van der Waals energy | -75.9 ± 8.3 |
| Electrostatic energy | -314.1 ± 43.8 |
| Desolvation energy | -20.6 ± 3.8 |
| Restraints violation energy | 32.7 ± 10.1 |
| Buried surface area | 2118.4 ± 66.6 |
| Z-score | 0.0 |

\*RMSD, root-mean-square deviation

### Preparation of a monomer mutant

The resulting dimer structure revealed that Trp17 and Trp24 were located at the dimer interface and interacted with arginine residues from the partner subunit (W17/R39 and W24/R16). Consequently, we hypothesized that replacing Trp with Arg would disrupt the cation-π interaction and introduce repulsion due to the positive charges. To create a monomeric mutant, we introduced three additional mutations, W24R, W17R, and W17/24R, in addition to the C78/157S mutation. However, mutations including W17R result in extensive precipitation during dialysis for refolding, rendering protein purification unfeasible. In contrast, VanX^C78/157S, W24R^ successfully produced the final product. Figure 9 illustrates the gel filtration chromatography elution profiles for VanX^C78/157S^ and VanX^C78/157S, W24R^, indicating the monomeric nature of VanX^C78/157S, W24R^. Notably, VanX^C78/157S, W24R^ exhibited greater instability than VanX^C78/157S^. We observed that salt precipitated proteins, particularly during the refolding step. Hence, prior to elution from the Ni-NTA column, we thoroughly washed the column with a urea-containing buffer to ensure the complete removal of guanidine hydrochloride from the elution fraction and increased the frequency of dialysis buffer changes during the early stages of refolding. Because of these adjustments, VanX^C78/157S, W24R^ did not precipitate during dialysis.

**Figure 9.**
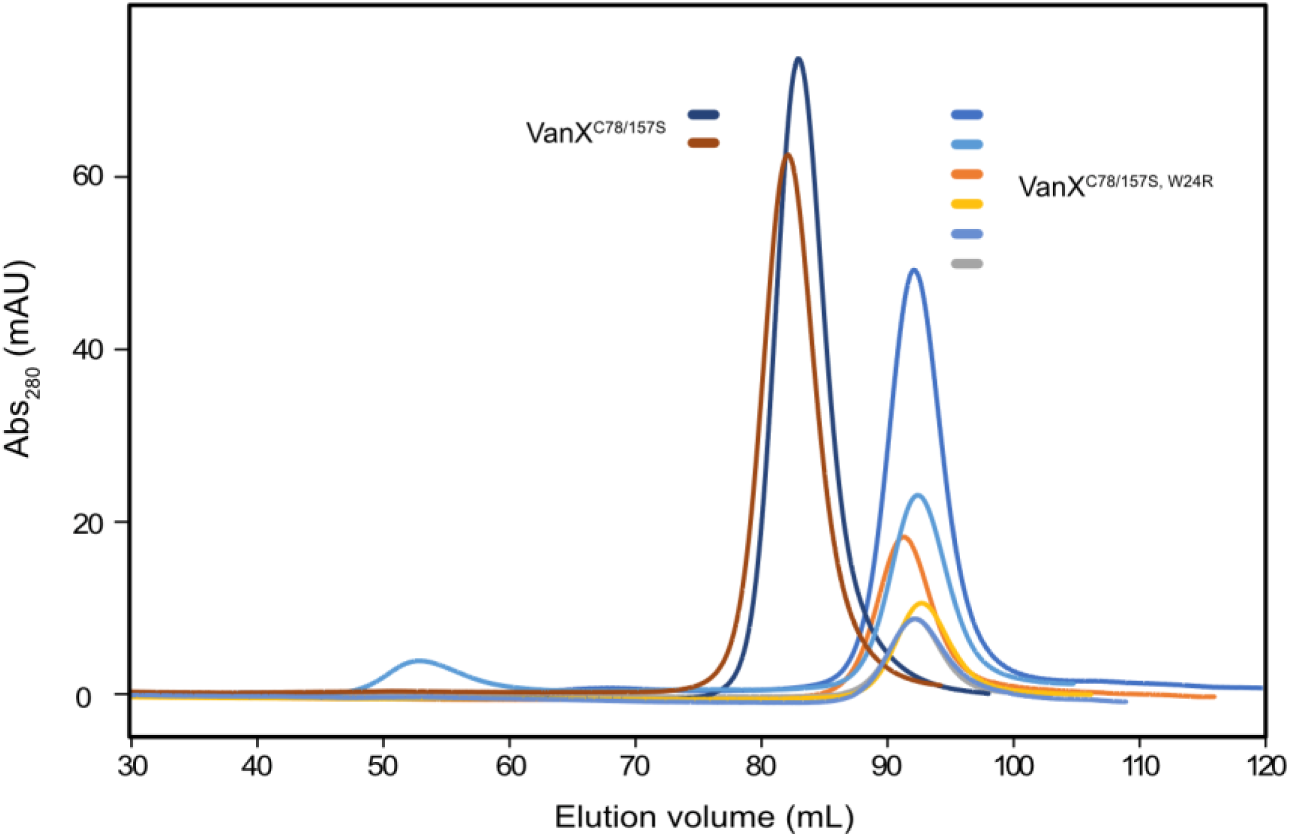
Overlay of the gel-filtration chromatography of VanX^C78/157S^ and VanX^C78/157S, W24R^. They were subjected to gel-filtration chromatography in the same way as described in the purification method. A total of two samples of VanX^C78/157S^ and six samples of VanX^C78/157S, W24R^ were separately analyzed on different days.

### Measurement of the monomer’s activity

The functional activity of the retained monomeric mutant was assessed. The substrate, D-Ala-D-Ala, contained two distinct methyl groups, each displaying distinct ^1^H NMR resonance frequencies, as shown in Figure 10 (a). Additionally, the methyl group present in the resultant product, D-Ala, exhibited a distinct ^1^H resonance value compared to that of either of the two methyl groups in the substrate. Using this feature, we measured a series of ^1^H 1D spectra to quantitatively determine enzymatic activity. NMR can measure activity in near real-time without the need for conventional methods, such as the modified cadmium ninhydrin reaction ^14,20,26^. The correlation between the molar ratio of the product and reaction duration is shown in Figure 10 (b). The graph illustrates that VanX sustained an almost constant rate of the forward reaction until nearly all substrates were converted into products, suggesting minimal occurrence of the reverse reaction. The *k*_cat_ values were determined from the slopes observed during the initial phase of the graph. Specifically, the calculated rates for the dimer and monomer were 8.6 and 4.7 ± 0.5 /s, respectively. The enzymatic activity of the dimer was lower than the previously reported *k*_cat_ value of 11 /s ^14^. This decrease could be attributed to the substitution of two cysteine residues with serine residues in the dimeric structure. The activity of the monomeric form was approximately half that of the dimer. As evidenced by the NMR 2D ^1^H/^15^N correlation spectra (see below), dissociation of the subunits introduced a minor perturbation to the overall structure, likely contributing to the observed decrease in enzymatic activity. Nonetheless, it was ascertained that the W24R mutation induced the monomerization of VanX while retaining at least half of its original activity.

**Figure 10.**
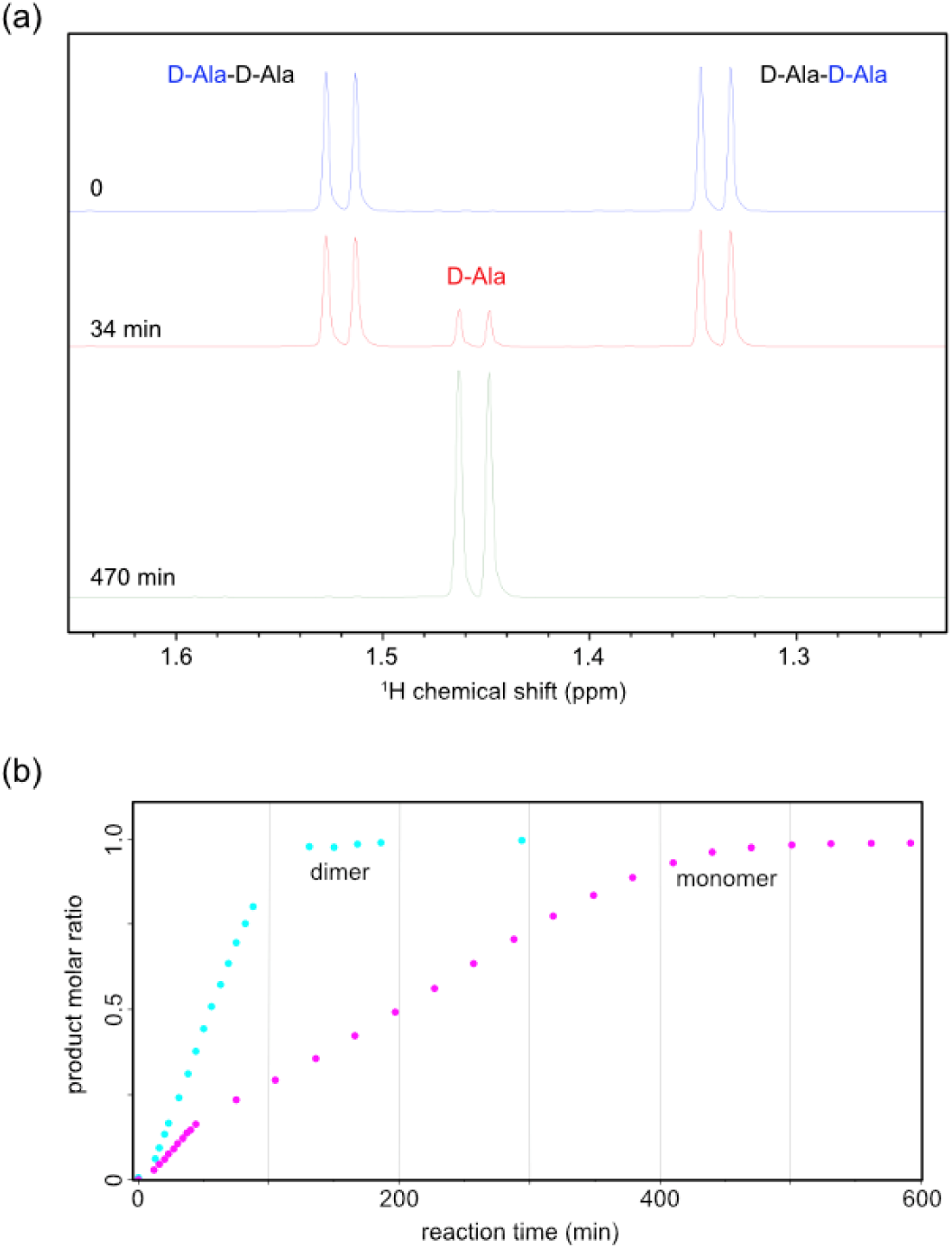
Activities of VanX monomer and dimer. (a) The ^1^H 1D NMR spectra of the substrate D-Ala-D-Ala and its product D-Ala are shown. The spectra represent measurements taken at 0, 34, and 470 min after the start of the reaction, catalyzed by VanX^C78/157S, W24R^. The two methyl groups were assigned based on a report ^27^. (b) The molar ratios of the product, D-Ala, were calculated from the peak intensities (integrals) of the ^1^H 1D spectra of VanX^C78/157S^ and VanX^C78/157S, W24R^ and plotted against the reaction time.

### Measurement of the monomer’s 2D NMR spectra

We purified the [^15^N, ^13^C]-VanX^C78/157S, W24R^ monomer and obtained its ^1^H-^15^N-HSQC spectrum. The superimposed spectra of the monomer and dimer are shown in Figure 11. Our analysis revealed that the NMR peaks in the monomer remained close to the resonance positions of the dimer, suggesting that monomerization had a minimal impact on the steric structure. Nonetheless, we observed slight shifts in many peaks, albeit of small magnitudes.

**Figure 11.**
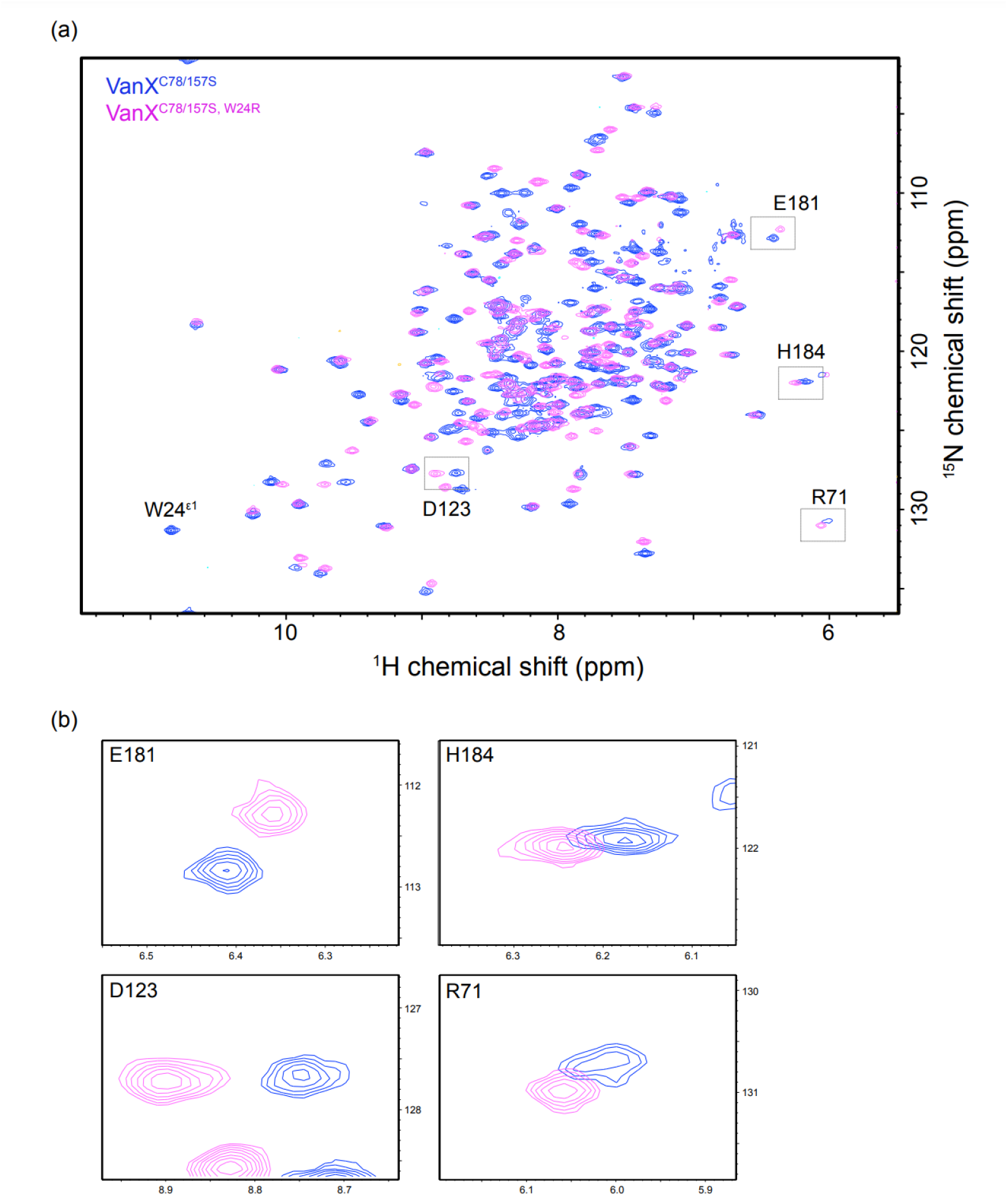
Overlaid 2D ^1^H-^15^N TROSY-HSQC spectra of 66 μM [^15^N, ^13^C]-VanX^C78/157S, W24R^ monomer (magenta) and 75 μM [^15^N]-VanX^C78/157S^ dimer (blue). The proteins were each dissolved in a 50 mM potassium phosphate buffer (pH 7.0) containing 30 μM ZnCl_2_ and 10% D_2_O. The spectra were measured at 303 K by a 500 MHz NMR spectrometer with 32 and 64 scans for the monomer and dimer, respectively. (a) Overall view. (b) Enlarged view of the peaks of residues around the active site.

## Discussion

### Antimicrobial resistance

A global survey in 2019 found that antimicrobial resistance caused more deaths than human immunodeficiency virus (HIV) or malaria ^28^. Of particular concern is MRSA, against which most antibiotics are ineffective. Vancomycin can suppress MRSA because its functional mechanism differs from that of β-lactam antibiotics, including penicillin. However, some bacterial strains develop resistance to vancomycin, which exacerbates problems, such as nosocomial infections. VanX is the key protein that mediates vancomycin resistance. The inhibition of VanX renders vancomycin-resistant bacteria susceptible to antibiotics, making it a valuable target for drug discovery. It is also anticipated that vancomycin sensitivity will increase when a VanX inhibitor is co-administered with D-Ala, a competitive inhibitor of VanA, a D-Ala-D-Lac ligase ^29^.

### Importance of NMR analyses

The development of inhibitors often relies on crystal structure analyses of the complexes formed between target proteins and candidate small molecules. Although crystal structures offer precise static atomic coordinates, they offer limited insights into comprehensive global or local site-specific dynamics. Temperature factors, although potentially informative about the dynamics, exhibit fluctuations within the crystal and may not accurately represent the conditions in solution. Isothermal titration calorimetry (ITC) and surface plasmon resonance (SPR) are frequently used for interaction analyses. However, these methods do not provide detailed insights into the site-specific local fluctuations of bound ligands or the interaction sites on proteins. In contrast, NMR spectroscopy can elucidate site-specific dynamics and atomic-level binding affinities. Consequently, the information derived from NMR, when combined with data from the crystal structures of complexes and other physicochemical analyses, is advantageous for enhancing candidate compounds toward more effective drug candidates with higher affinity.

### VanX as a homodimer

The crystal structure of VanX has been shown to be a hexamer ^6^. Although Brandt et al ^30^ judged that VanX assumed a dimeric form based on MALDI-TOF mass spectrometry, their spectrum also exhibited peaks corresponding to monomeric and tetrameric species, albeit at low levels. Wu *et al.* ^14^ estimated the molecular mass of VanX to be 38 kDa using gel filtration chromatography. Our gel filtration experiments also showed an apparent molecular mass slightly smaller than the 43 kDa marker. Although we cannot rule out the possibility of a non-specific interaction between VanX and the resin, these results suggest that at such a low concentration, as in a gel-filtration column, VanX may exist in equilibrium between the major dimeric (46 kDa) and minor monomeric (23 kDa) states. Indeed, our finding that the substitution of a single residue, Trp24, leads to monomerization is consistent with this hypothesis. Nonetheless, we have so far measured hundreds of NMR 2D ^1^H-^15^N correlation spectra across a range of NMR-detectable subunit concentrations from 20 to 200 μM and observed no discernible concentration-dependent shifts in peak positions. If an exchange equilibrium exists in the interactions among the subunits, the peaks would either assume resonances among those of the monomeric, dimeric, and tetrameric states (fast exchange mode) or concurrently display resonances of these states (slow exchange mode). Additionally, if an intermediate exchange mode is present on the NMR timescale of microseconds to milliseconds, the NMR peaks would generally tend to broaden, making observations difficult. However, no such phenomena suggesting the hypothesis were observed in our spectra. Thus, under NMR measurement conditions, it is reasonable to conclude that VanX predominantly adopts a homodimeric state.

### Contribution of RDC to modeling multimer structures

We prepared [^2^H, ^15^N, ^13^C]-VanX^C78/157S^ for the RDC measurements. This labeling pattern was chosen because the 2D ^1^H-^15^N correlation spectrum of non-deuterated VanX showed broadening, particularly of the ^15^N-anti-TROSY peaks, making it difficult to obtain accurate RDC values. Even after deuteration, we encountered severe peak overlaps, which prevented us from obtaining precise RDC values for certain residues. To address this problem, we generated a 3D spectrum by acquiring ^1^H^N^-TROSY and anti-TROSY peaks in the TROSY-HNCO experiment. However, because the sensitivity of the 3D spectrum was relatively low, we supplemented it with a 2D spectrum to obtain the RDC values from either of the spectra. Despite our efforts, as shown in Figure 7, we did not achieve the best correlation between the measured and back-calculated RDC values for the subunit (Q-factor: 45.9%). Chen *et al.* have described that structures with resolutions of 2–3 and 1.5 Å commonly exhibit Q-factors of 40 and 20%, respectively ^31^. There are several possible reasons for this discrepancy. One possibility is that the amide groups of flexible residues have smaller absolute RDC values, allowing us to exclude data from the loop regions exposed on the molecular surface. Additionally, because the crystal structure lacked the coordinates of ^1^H atoms, we had to calculate them using a tool that might not perfectly reflect the actual ^1^H positions. However, as VanX is a homodimer, one of the three principal axis of the molecular alignment tensor should be aligned with a two-fold rotational symmetry axes ^32^. This special feature notably enhanced the accuracy of the relative orientational analysis of identical subunits using RDC. Another factor to consider is the structural distortion caused by crystal packing.

### Monomerization by a single mutation

A comparison of the NMR spectra of the dimer VanX^C78/157S^ and the monomer VanX^C78/157S, W24R^ revealed that the overall peak positions were similar (Figure 11); however, residues showing shifted peaks were observed throughout the entire subunit. Because changes in the amide ^1^H chemical shifts are sensitive to the hydrogen bond distances ^33^, even though the peak displacements may appear significant, it is anticipated that the shifts in residues within the stereochemical structure are minimal. In other words, the effects of transitioning from dimer to monomer are distributed throughout the entire molecule, causing slight movements of individual atoms that lead to an overall adjustment of hydrogen bonding. This is believed to be the cause of the 50% decrease in activity. The monomer exposes the hydrophobic subunit interface, which was originally hidden in the dimers. Although the Arg mutation introduced a positive charge, it created nonspecific hydrophobic interactions. This was evident from the fact that high salt concentrations destabilized the monomer during the purification process. However, in the future, when using NMR for drug discovery, including screening, half of the molecular weight will provide a more sensitive spectrum than the dimer. Specifically, the analyses of proteins with molecular mass greater than 50 kDa often require deuteration, which interferes with the assignment and observation of ^1^H side chains. In contrast, the VanX monomer does not require deuteration, which is a remarkable advantage. We are currently investigating additives such as glycerol, arginine, and sulfobetaine ^34^ to increase the stability of the monomer, which will enable more precise side chain assignments in regions containing the active site. We believe that most assignments in the monomer can be applied to the corresponding dimer assignments because ^1^H/^13^C does not participate in hydrogen bonding, and the changes in chemical shifts due to subunit splitting are expected to be smaller than those of ^1^H/^15^N amide groups. This maximizes the use of NMR for the development of inhibitors targeting VanX ^35^.

### Summary

The crystal structure of VanX has been shown to be a linear hexamer ^6^. However, we confirmed that VanX formed a stable homodimer in solution and determined its structural model. Specifically, we identified the interface between the subunits using the NMR TCS and NOE methods and constrained the relative orientation of the subunits using RDC. These results closely match the B/C subunits in the crystal structure (1r44). Furthermore, by substituting Trp24 at the interface with Arg, a monomeric mutant that retains half of its activity was successfully generated. The dimer with two cysteines replaced by serines showed enhanced stability compared to the wild-type protein. The amide group assignment data obtained will be valuable for future NMR-based inhibitor development. The monomeric mutant, which has a low molecular weight, does not require ^2^H labeling and allows for the observation of side chains through NMR. This has the potential to aid in the development of drugs to combat vancomycin-resistant bacteria.

## Experimental Procedures

### Expression and purification of isotopically labeled VanX

Genes encoding VanX (C78/157S mutants with and without W24R) were constructed using the pET-47b plasmid vector (Novagen). The HRV3C recognition sequence and a linker to improve the digestion efficiency were inserted between the (His)_6_-tag and the *vanx* sequence. This resulted in the VanX polypeptide with the sequence GPGMGGSG added to its N-terminus after digestion with GST-fused HRV3C. *E. coli* strain BL21(DE3) pLysS, transformed with either plasmid, was cultured in an M9 minimal medium containing 20 µg/mL kanamycin, 1.0 g/L [99% ^15^N]-ammonium chloride (^15^NH_4_Cl), and 2.0 g/L [99% ^13^C_6_]-D-glucose at 37°C. For the expression of perdeuterated VanX, an M9 medium was prepared with 99% D_2_O containing 2.0 g/L [>97% ^2^H_7_, >98% ^13^C_6_]-D-glucose (Cambridge Isotope Laboratories, Inc.). When the optical density at 600 nm (OD_600_) reached 0.4, 0.5 mM isopropyl β-D-1-thiogalactopyranoside (IPTG) was added for induction. After three hours, cells were collected by centrifugation. The cells were suspended in 20 mM Tris-HCl buffer (pH 8.0) containing 6 M guanidine hydrochloride, and disrupted by sonication. For VanX^C78/157S, W24R^ and VanX^C78/157S, W17R^ mutants, sonication was employed in a guanidine hydrochloride-free buffer because almost all expressed proteins were found within the inclusion bodies, and the gathered precipitates were solubilized by adding a buffer containing guanidine hydrochloride. This approach leads to a reduction in impurities. The supernatant was subjected to Ni^2+^-immobilized affinity chromatography (Chelating Sepharose Fast Flow; Cytiva). The resin was washed with the same buffer and with another buffer containing 8 M urea instead of guanidine hydrochloride, both containing 10 mM imidazole. Unfolded (His)_6_-tagged VanX was eluted with 20 mM Tris-HCl buffer (pH 8.0) containing 8 M urea and 300 mM imidazole. The eluate, diluted to < 1.5 μM in a dialysis tube with a molecular weight cut-off of 12,000–14,000, was dialyzed overnight at 4°C against 100 times the amount of 20 mM Tris-HCl buffer (pH 7.5) containing 1 mM ethylenediaminetetraacetic acid (EDTA). To VanX refolded by dialysis, GST-fused HRV3C was added (1/30 wt/wt) overnight at 4°C to cleave the (His)_6_-tag. Since the isoelectric points of Hrv3v and GST are 8.46 and 4–5, respectively, we avoided using a dialysis buffer at pH 8. Finally, the sample was subjected to size-exclusion chromatography (SEC) using a HiLoad 16/60 Superdex 200 pg column (Cytiva) with 20 mM Tris-HCl running buffer (pH 8.0) containing 150 mM NaCl and 30 µM ZnCl_2_ at a flow rate of 1 mL/min. In the purification of VanX^C78/157S, W24R^, 5% glycerol was added during concentration and SEC to prevent precipitation. The eluted protein was concentrated using a centrifugal filter unit (Amicon Ultra, 10 or 30 kDa cut-off, Merck), and the solvent was exchanged with 50 mM potassium phosphate buffer (pH 7.0) containing 30 µM ZnCl_2_ and 10% D_2_O for subsequent NMR measurements.

### Amino acid-selective unlabeling

*E. coli* cells were cultured in 100 mL of Luria-Bertani (LB) medium. When an OD_600_ reached 0.3, the medium was centrifuged, and the pellet cells were transferred to 500 mL of M9 minimal medium containing 1.0 g/L ^15^NH_4_Cl, 2.0 g/L D-glucose, and 1.0 g/L of one of the desired unlabeled amino acids. Expression was induced by adding 1 mM IPTG at an OD_600_ of 0.3. Three hours later, cells were collected by centrifugation. Each sample was dissolved at a subunit concentration of 35–75 µM in 50 mM potassium phosphate buffer (pH 7.0) or 20 mM sodium acetate buffer (pH 4.0), both containing 30 µM ZnCl_2_ and 10% D_2_O. For each sample, a ^1^H-^15^N TROSY-HSQC spectrum was measured at 303 K with a total of 2,048 data points for ^1^H and 256 data points for ^15^N using a ^1^H resonance frequency of 500.13 MHz for 8 h.

### Main-chain assignment of the dimer

The NMR spectra for the backbone assignment of the dimer were acquired at 303 K using a Bruker BioSpin Avance III HD spectrometer equipped with a TCI triple-resonance cryogenic probe with a ^1^H resonance frequency of 800.23 MHz. Two-dimensional (2D) ^1^H-^15^N heteronuclear single-quantum correlation (HSQC), three-dimensional (3D) HNCACB, HN(CO)CACB, HNCA, HN(CO)CA, HNCO, and HN(CA)CO spectra were acquired using 100 µM [^2^H, ^15^N, ^13^C]-VanX^C78/157S^. All experiments utilized the ^1^H/^15^N TROSY technique ^36^, and data in the ^13^C dimension were sampled using the States-TPPI method. The WATERGATE method was employed to suppress the water signal ^37^. Non-uniform sampling was applied to the ^13^C and ^15^N dimensions. The main-chain assignments were also made using 100 µM [^2^H, ^15^N, ^13^C]-VanX^C78/157S^ dissolved in 20 mM deuterated sodium acetate buffer (pH 4.0) containing 30 µM ZnCl_2_ and 10% D_2_O. All NMR data were processed using NMRPipe ^38^, and the spectra were analyzed using NMRFAM-Sparky ^39^.

### TCS experiment

TCS experiments were conducted using a sample containing 0.2 mM [^15^N, ^2^H]-VanX^C78/157S^ and 0.2 mM [^13^C, ^1^H]-VanX^C78/157S^ in a 50 mM potassium phosphate buffer (pH 7.0) containing 30 µM ZnCl_2_ and 10% D_2_O. The two types of differently labeled VanX were mixed before refolding. Because of the low sensitivity of TCS in a solvent with a composition of 10% H_2_O and 90% D_2_O, we used a solvent composition of 90% H_2_O and 10% D_2_O. The experiments were performed at 303 K using an NMR spectrometer (Bruker BioSpin Avance III HD) equipped with a TCI triple-resonance cryogenic probe operating at a ^1^H resonance frequency of 500.13 MHz. The experimental setup followed the procedure described by Takahashi et al. ^40^. A relaxation delay of 2.0 s was applied before ^1^H saturation, which was achieved by repeating a delay of 10 ms and a 10 ms-long RSNOB pulse with a phase x/-x, a maximum amplitude of 100 or 200 Hz, and an excitation center at 1.0 ppm. The saturation time lasted for 0.5, 1.0, or 2.0 s with the 100 Hz amplitude or 1.0 s with the 200 Hz amplitude, followed by a ^1^H-^15^N TROSY-HSQC sequence incorporating a 3-9-19 WATERGATE scheme. To obtain a reference spectrum, interleaved acquisition was employed by shifting the excitation center of the saturation pulses to 30,000 Hz off-resonance. The spectral widths were set to 24 ppm (2,048 data points) for the ^1^H dimension and 35 ppm (300 data points) for the ^15^N dimension in each spectrum. Each free induction decay (FID) consisted of 32 scans. The carrier frequencies were positioned at 4.7 and 119 ppm for the ^1^H and ^15^N dimensions, respectively.

### Measurement of RDC constants

For the analysis of the ^1^H/^15^N residual dipolar coupling (RDC) constants, the coat protein of *Pf1* filamentous bacteriophage (*Pf1*-LP11-92 in a 10 mM potassium phosphate buffer (pH 7.6) containing 2 mM MgCl_2_ and 0.05% NaN_3_, ASLA Biotech) was added as an alignment medium at a concentration of 27 mg/mL to a solution of 154 µM [^2^H, ^15^N, ^13^C]-VanX^C78/157S^ in a 50 mM potassium phosphate buffer (pH 7.0) containing 30 µM ZnCl_2_, 10% D_2_O, and 0.02% NaN_3_. The spectra (^1^H-^15^N TROSY/anti-TROSY-based HSQC) were measured at 303 K using an 800 MHz NMR spectrometer. The pulse program was modified from a Bruker’s standard pulse program (trosyf3gpphsi19.2) to acquire ^15^N-TROSY and anti-TROSY peaks in an interleaved manner. The acquisition times for the ^1^H and ^15^N dimensions were 64 and 45 ms, respectively. Using NMRPipe ^38^, the ^15^N dimension (35 ppm range) was extended to 4,096 points using zero-filling. For the 3D spectral measurements, the TROSY and anti-TROSY peaks split along the ^1^H dimension were acquired in an interleaved manner at 303 K using a 500 MHz NMR spectrometer. This was achieved by modifying a Bruker’s standard TROSY-HNCO pulse program (trhncogp3d). The acquisition times for the ^1^H, ^15^N, and ^13^C’ dimensions were set to 79, 21, and 25 ms, respectively, with the adoption of 70% non-uniform sampling. The time-domain data were processed using SMILE in NMRPipe ^41^, and the peaks were picked using NMRFAM-Sparky ^39^. The RDC data were analyzed using the coordinates of subunits A and B in the PDB (1r44), to which the amide hydrogens were added, using software in the PDB Utilities Server of the Bax group (NIH) developed by Dr. F. Delaglio (https://spin.niddk.nih.gov/bax/nmrserver/index.html).

### Calculation of the dimer conformation

The atomic coordinates of subunit B were retrieved from PDB:1r44 and duplicated as the coordinates of homodimers for input into HADDOCK 2.4 ^42,43^. The N- and C-termini were positively and negatively charged, respectively. During the initial stage (it0) of the structural calculations, coarse-grain modeling was omitted, and the original positions were left unconstrained. Rather than explicitly specifying the passive residues, ambiguous restraints based on the TCS experimental results were applied. Fifty percent of the ambiguous restraints were randomly excluded according to the procedure described in the “Random AIR definition (*ab-initio* mode)” (https://www.bonvinlab.org/software/haddock2.4/airs/). The distance constraint between Trp24 and Leu134 was applied as the active residues on the basis of the observed intersubunit NOE. During the calculation process, nonpolar hydrogen atoms were excluded using the default settings. *C*_2_ symmetry constraints with a force constant of 10.0 were imposed. The RDC values were transformed to the input format on the basis of the procedure described in the webpage “HADDOCK2.4 manual - Residual Dipolar Couplings” (https://www.bonvinlab.org/software/haddock2.4/RDC/). The RDC type selected was SANI, and the initial alignment tensor values (*D*a and *D*r) were estimated using Pales ^25^. Other parameters and settings were obtained from the defaults.

### Activity measurements

The activities of VanX^C78/157S^ and VanX^C78/157S, W24R^ were assessed using NMR in the following manner. Initially, a substrate solution was prepared by dissolving 10 mM D-Ala-D-Ala in 590 µL of buffer, composed of 50 mM potassium phosphate (pH 7.0), 30 µM ZnCl_2_, and 10% D_2_O. The 1D ^1^H NMR spectrum of the substrate was recorded as a reference at time zero at 298 K using a 500 MHz NMR spectrometer. A 1D pulse sequence with 3-9-19 WATERGATE was used to suppress the water signal. The ^1^H excitation pulse length (p1) was set to 3 μs. The inter-scan delay was set to 5 s. Each 1D experiment took 3.4 min. Subsequently, VanX^C78/157S^ or VanX^C78/157S, W24R^ was added to the substrate solution, resulting in the final subunit concentrations of 0.19 and 0.16 µM, respectively. Immediately after the addition, a 1D ^1^H NMR spectrum was acquired. Further measurements were performed at predefined time intervals to monitor the reactions. During the measurements, alterations in the peaks corresponding to the methyl groups of D-Ala-D-Ala and D-Ala were observed. Upon integration and normalization of the peak areas, we estimated the molar ratios of the products, which were then plotted against reaction time. By analyzing the obtained graph, we calculated the initial reaction rates, denoted as *k*_cat_ (number of D-Ala molecules produced by one subunit per second).

## Author Contributions

T.K., A.N., S.A., T.Y., H.A., and T.I. designed the experiments; T.T., C.T., M.N., M.H., Y.Y., H.S., E.C., and K.A. prepared the samples and performed the experiments; and T.I. wrote the paper.

## Acknowledgments

T.I. expresses gratitude to Ms. Yoko Motoda, Dr. Tomoki Matsuda, Dr. Yutaka Kuroda, Dr. Takahisa Nakai, and Dr. Haruki Nakamura for their invaluable assistance and discussions. This study used a cloud system, NMRbox: National Center for Biomolecular NMR Data Processing and Analysis, BTRR.

## Additional data

The chemical shifts of the main chain at pH 4 and 7 have been registered under the accession number 52158 in the Biological Magnetic Resonance Data Bank (BMRB).

---

**Figure S1.**
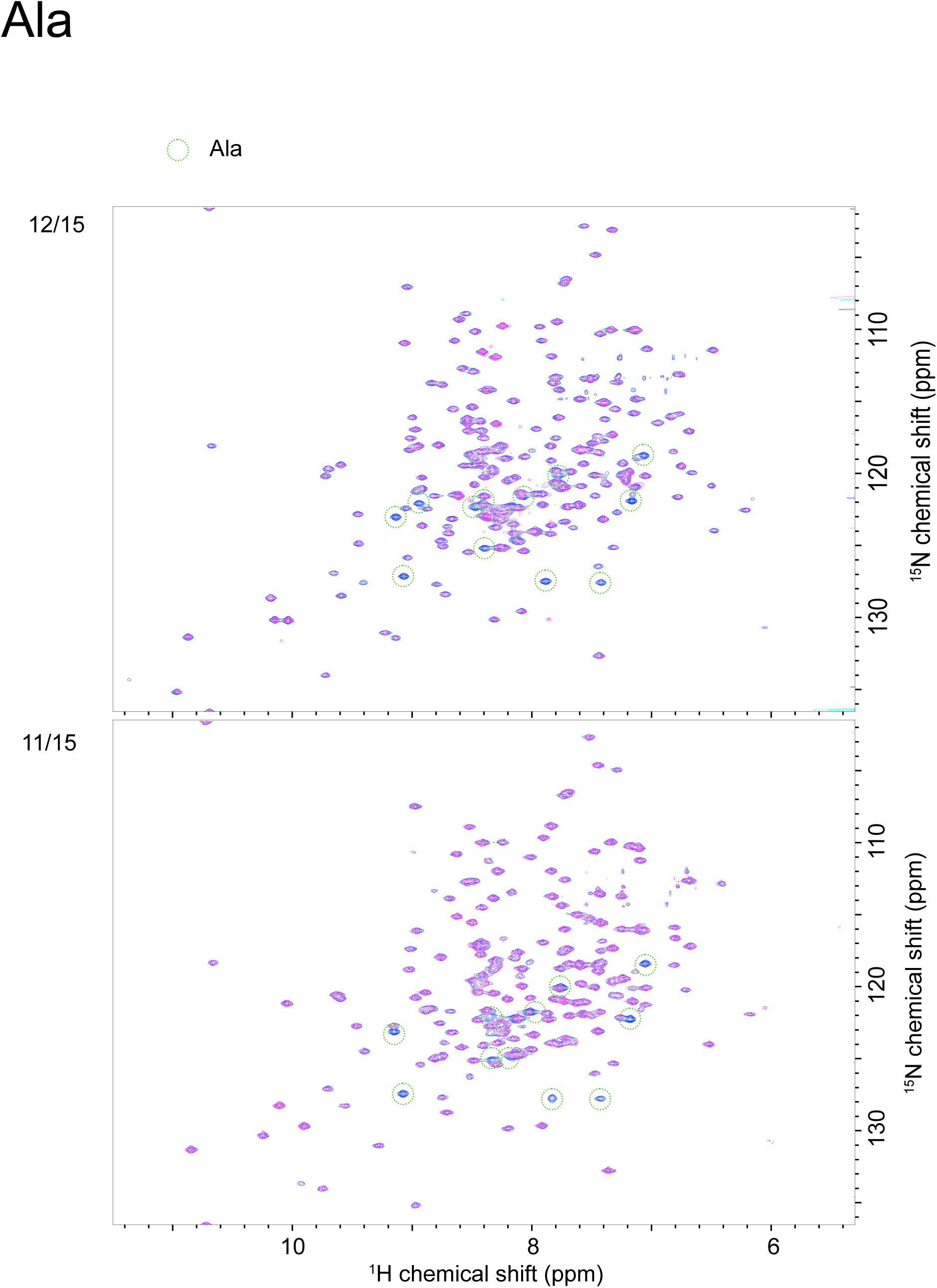
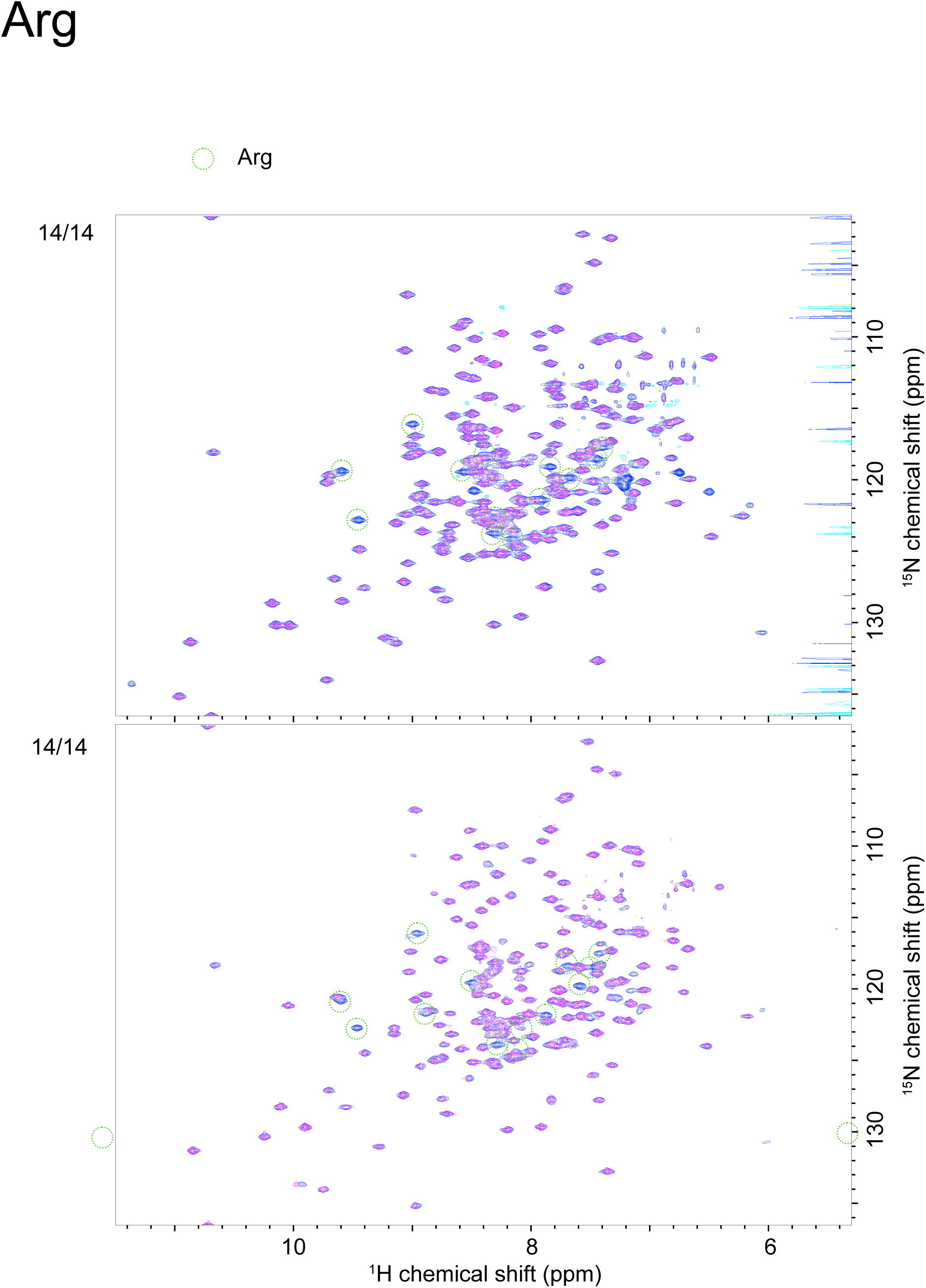
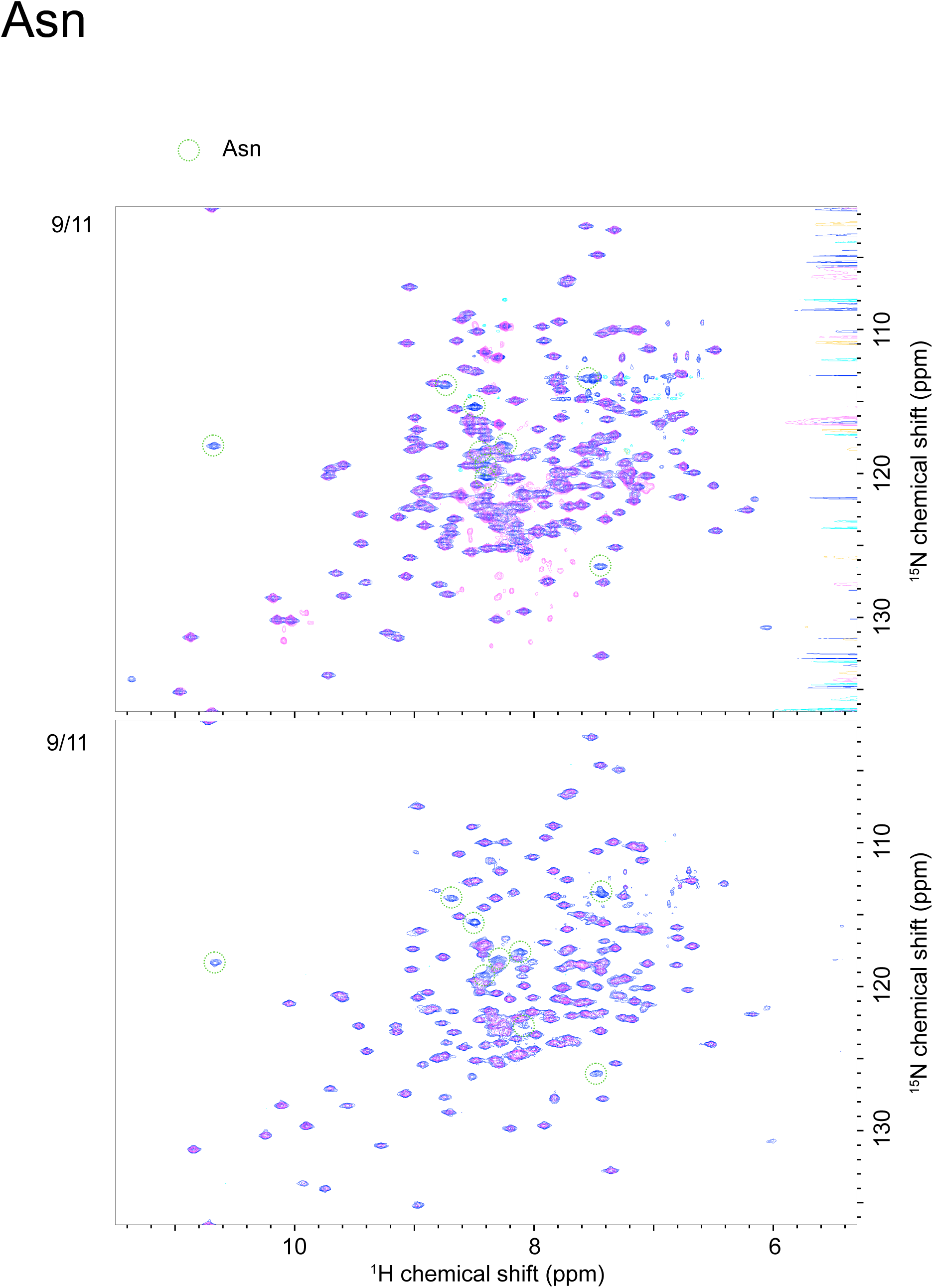
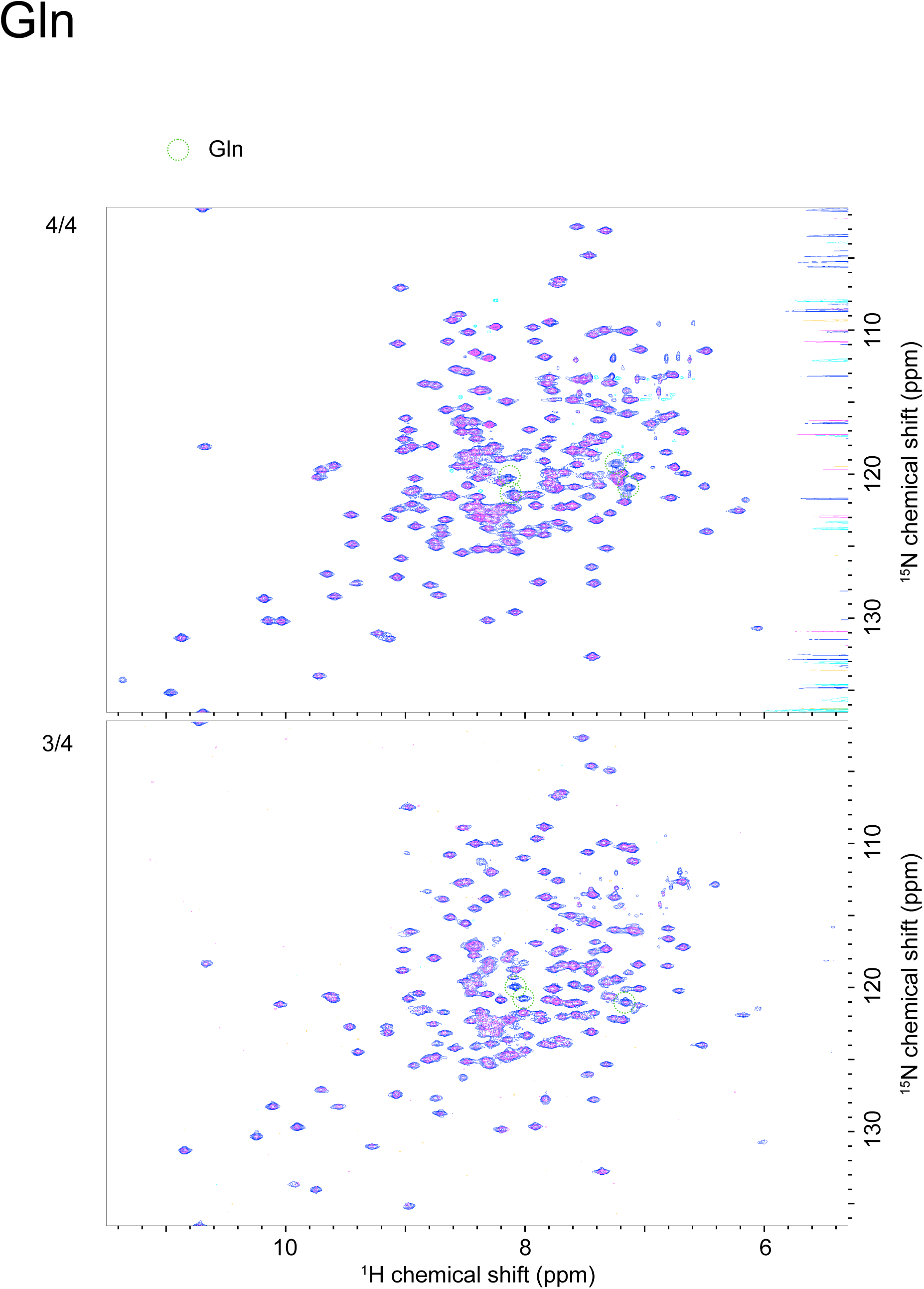
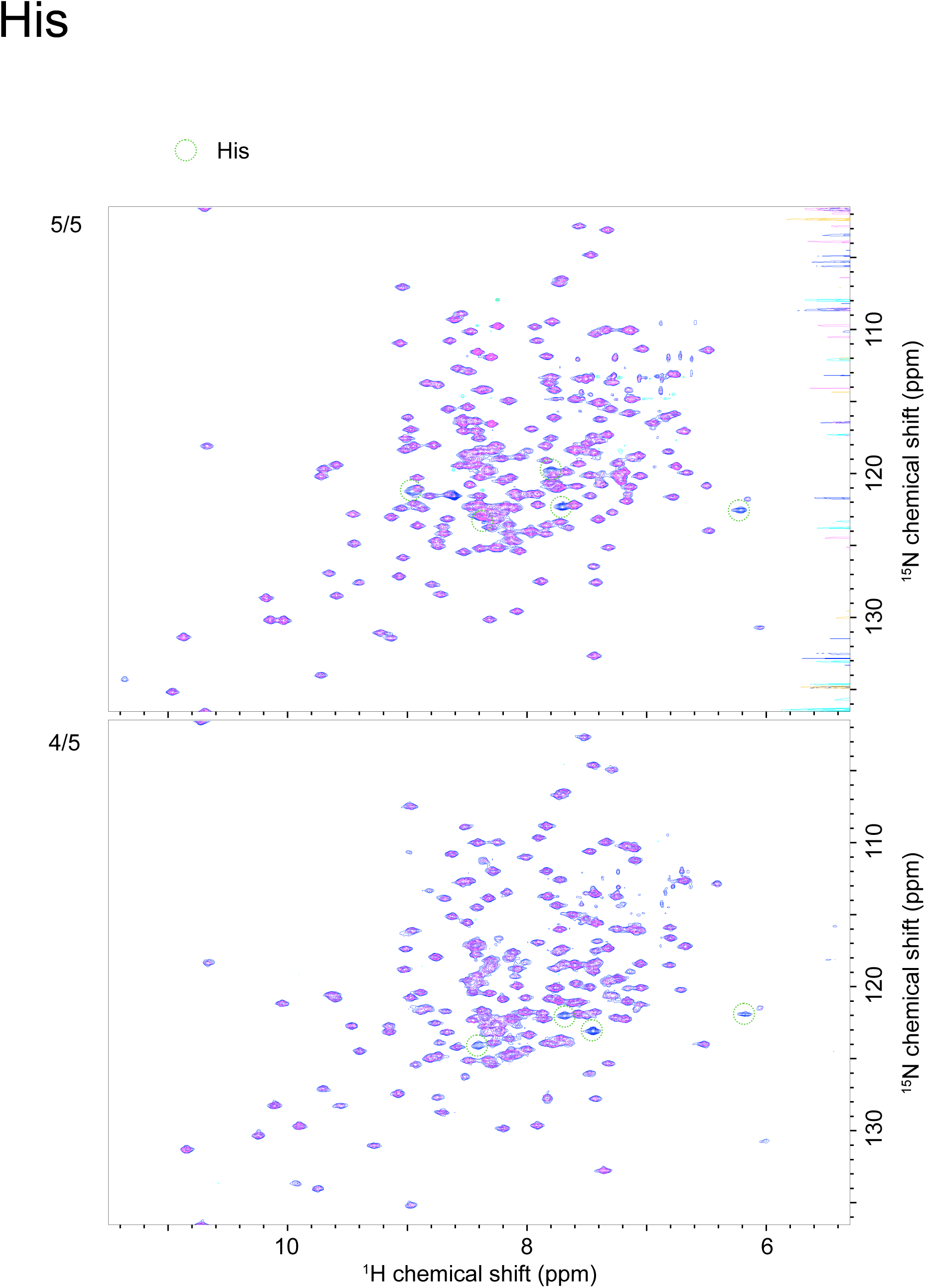
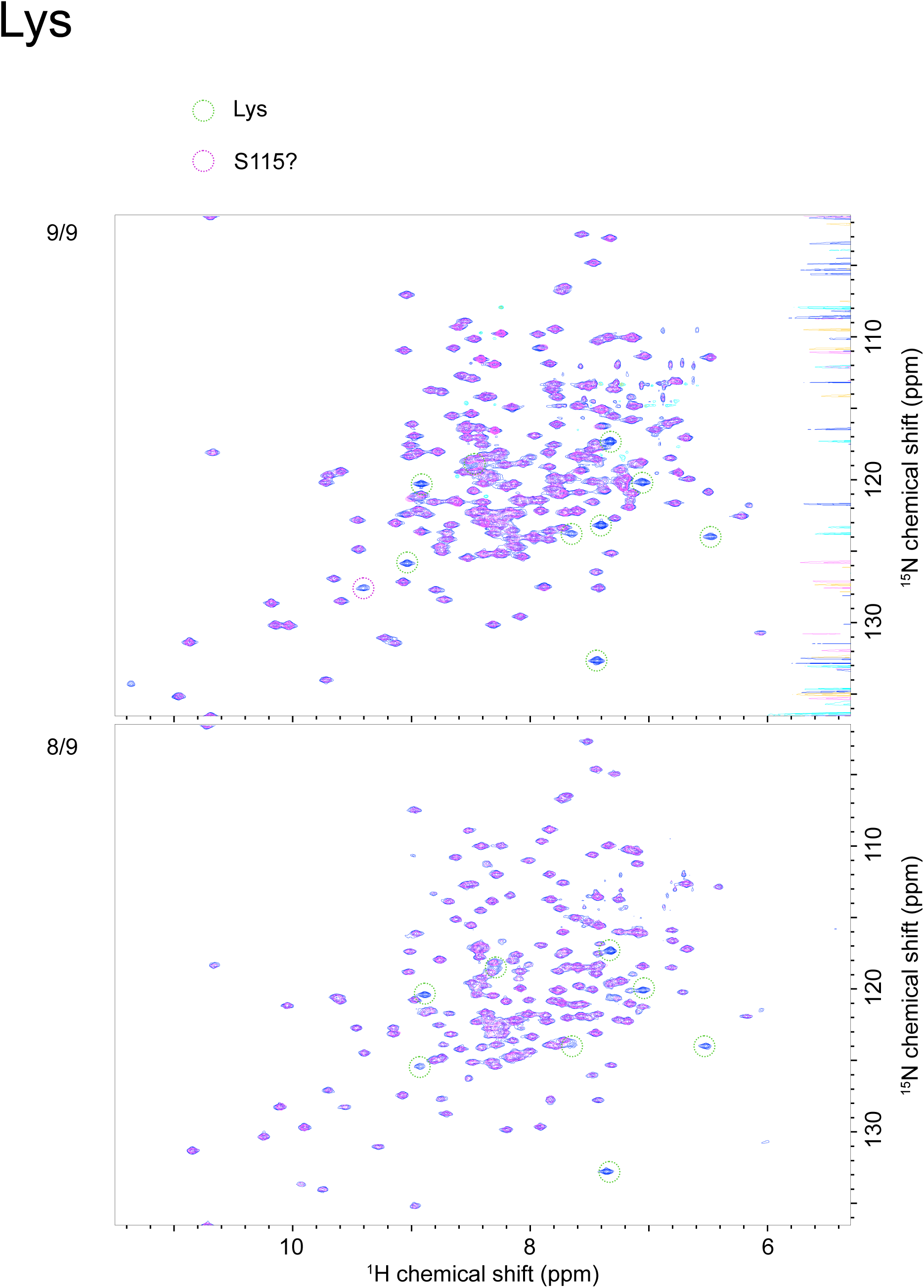
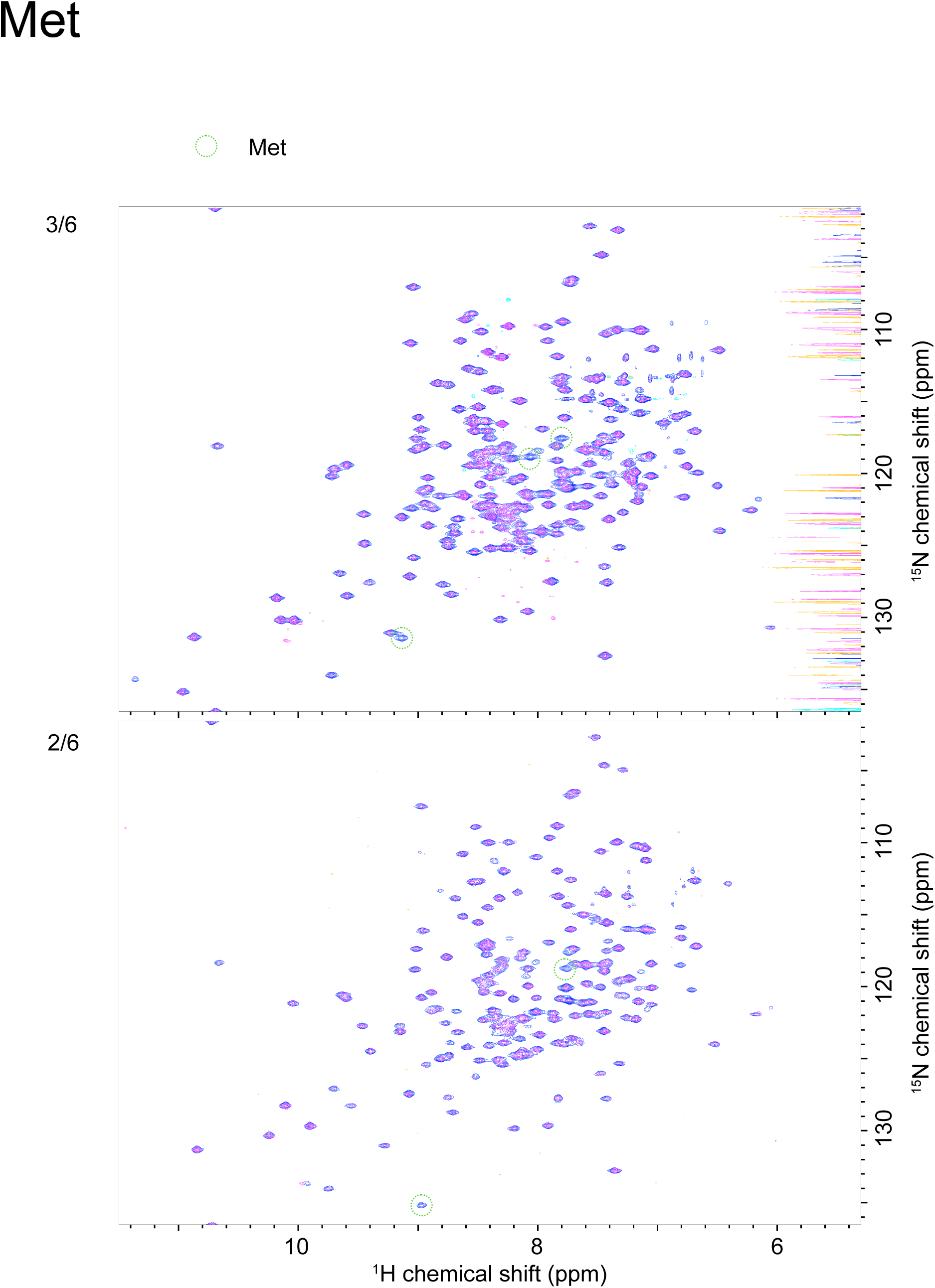
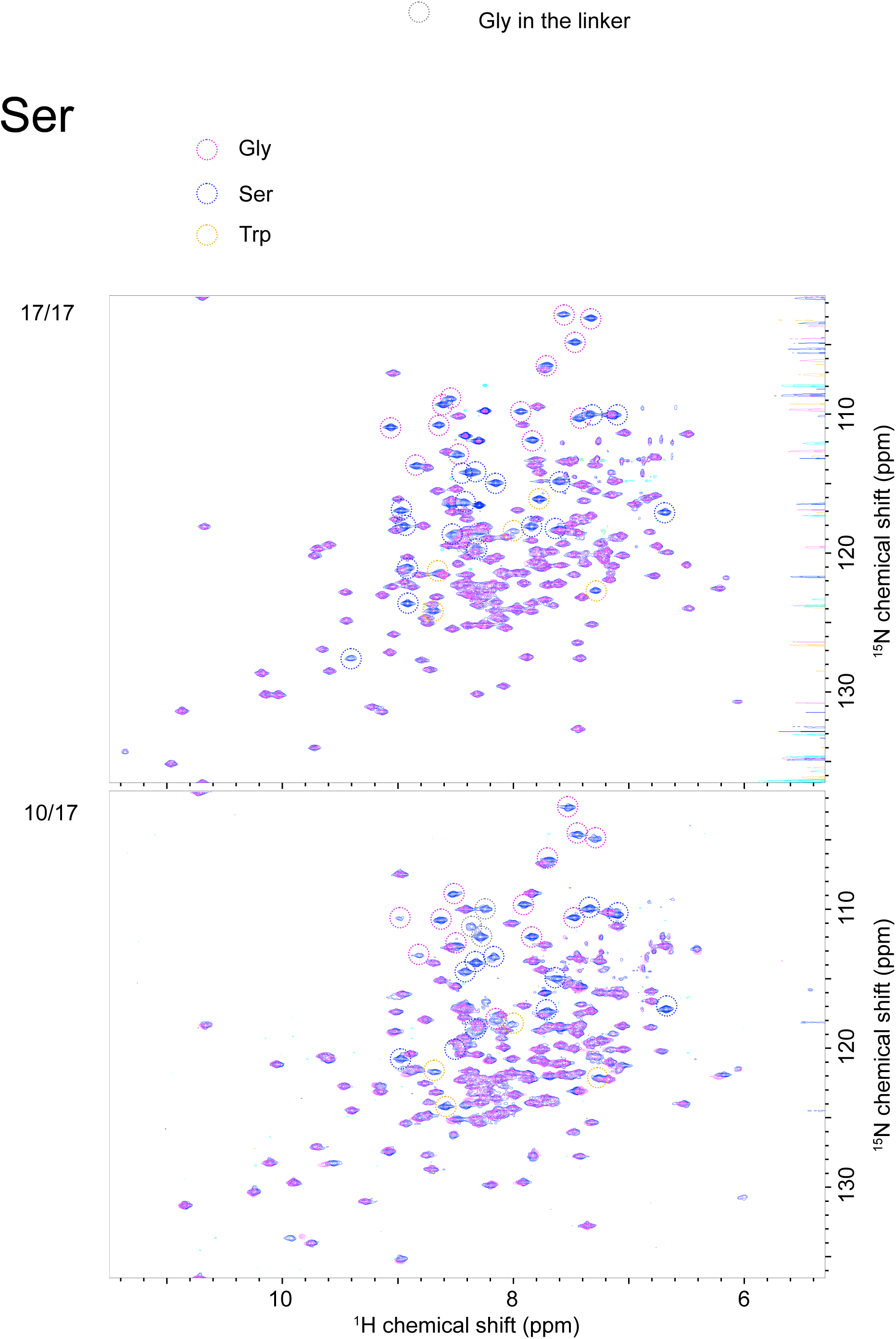
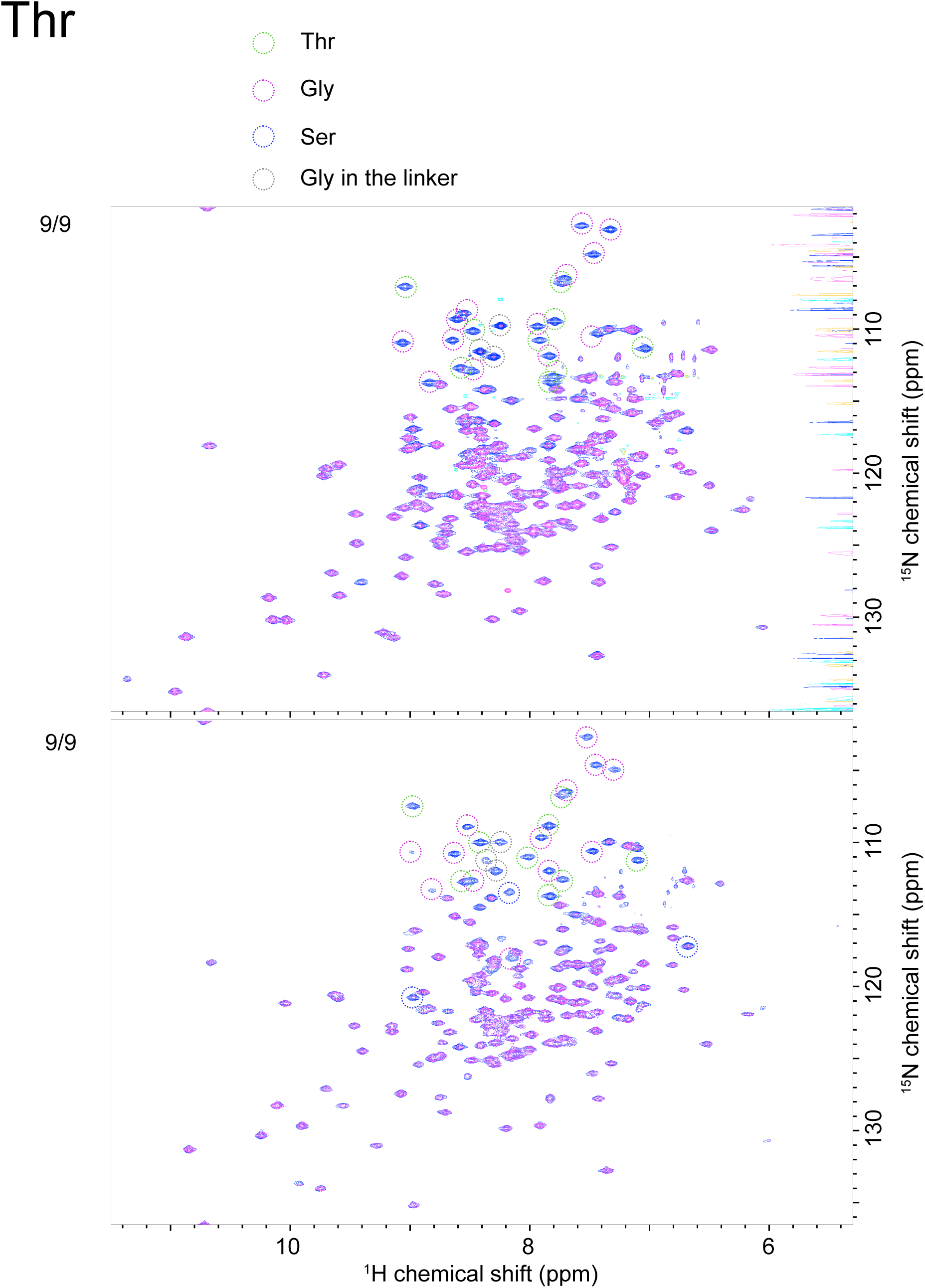
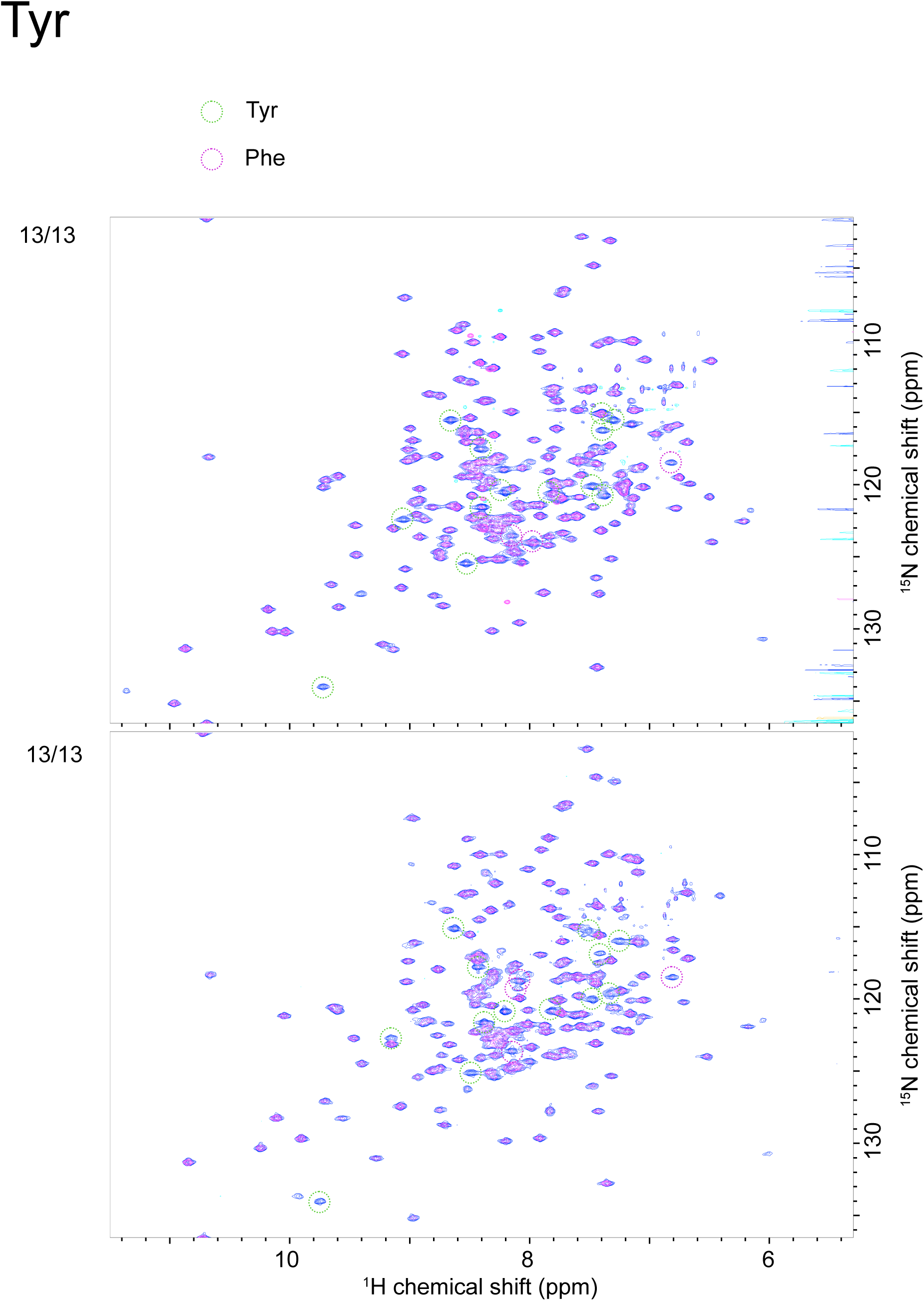
Overlay of 2D ^1^H-^15^N TROSY-HSQC spectra of reference VanX (blue) and VanX with each specific amino acid unlabeled (magenta). The pH 4 and pH 7 spectra are on the upper and lower sides, respectively. Green circles indicate unlabeled target amino acids. Circles of other colors indicate amino acids for which the ^15^N labeling ratio was reduced, attributable to metabolic scrambling during growth in *E. coli*. The fraction *n*/*m*, written on the left side of each spectrum, indicates that *n* out of the expected *m* peaks were actually broadened.

**Figure S2.**
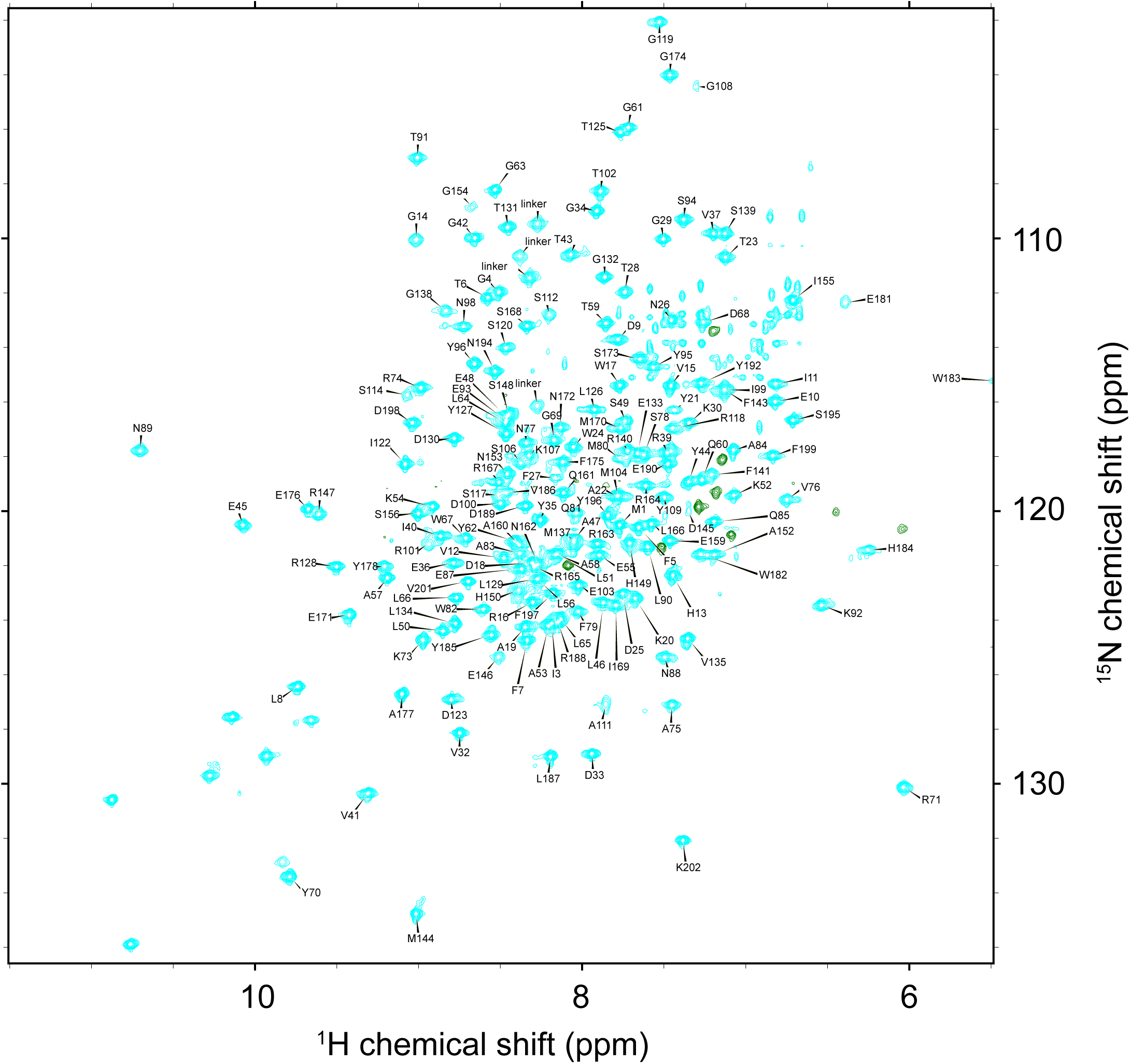
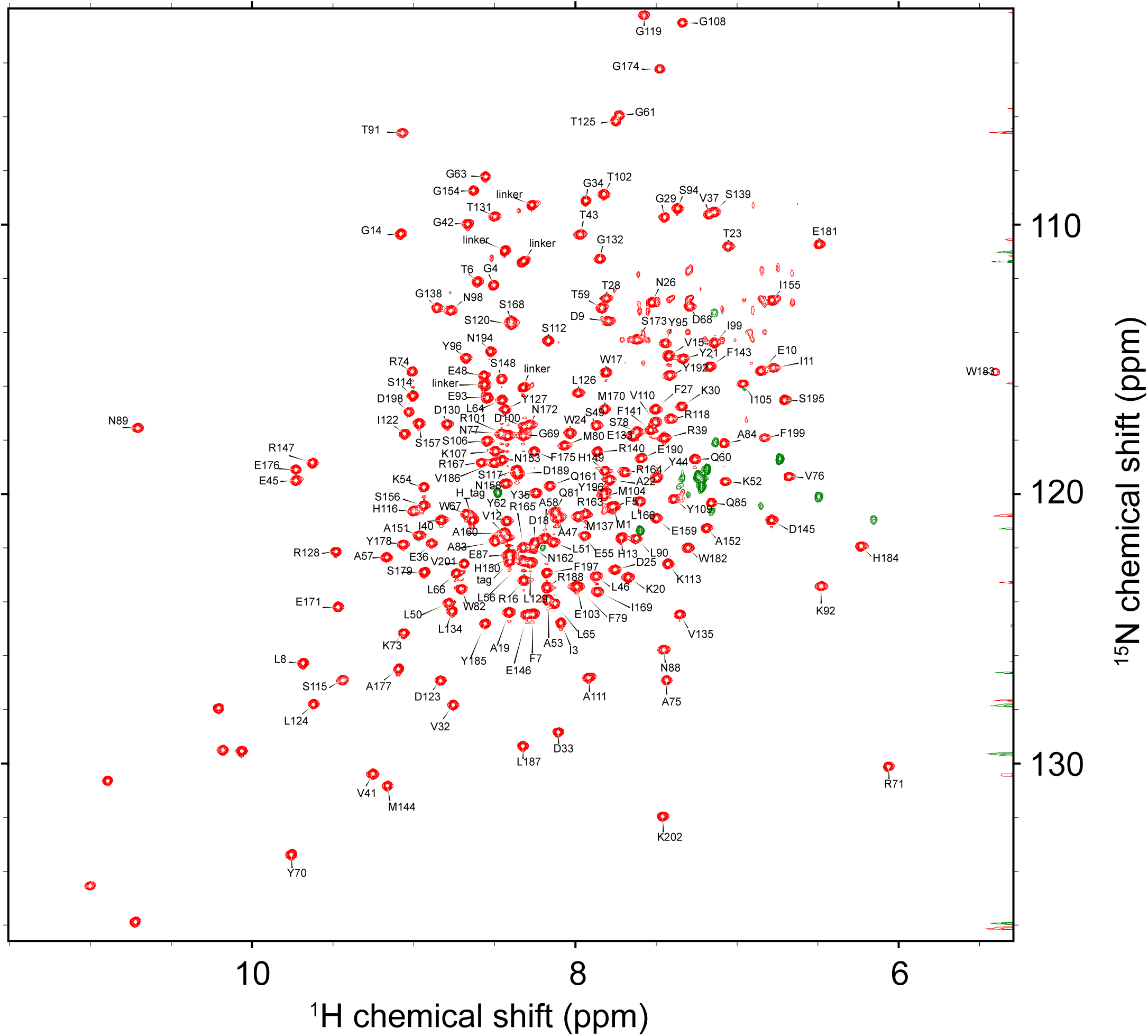
^1^H-^15^N TROSY-HSQC spectra of [^2^H, ^15^N, ^13^C]-VanX^C78/157S^ at pH 7 (cyan) and 4 (red). Peaks were annotated and assigned to the corresponding main-chain amide groups.

**Figure S3.**
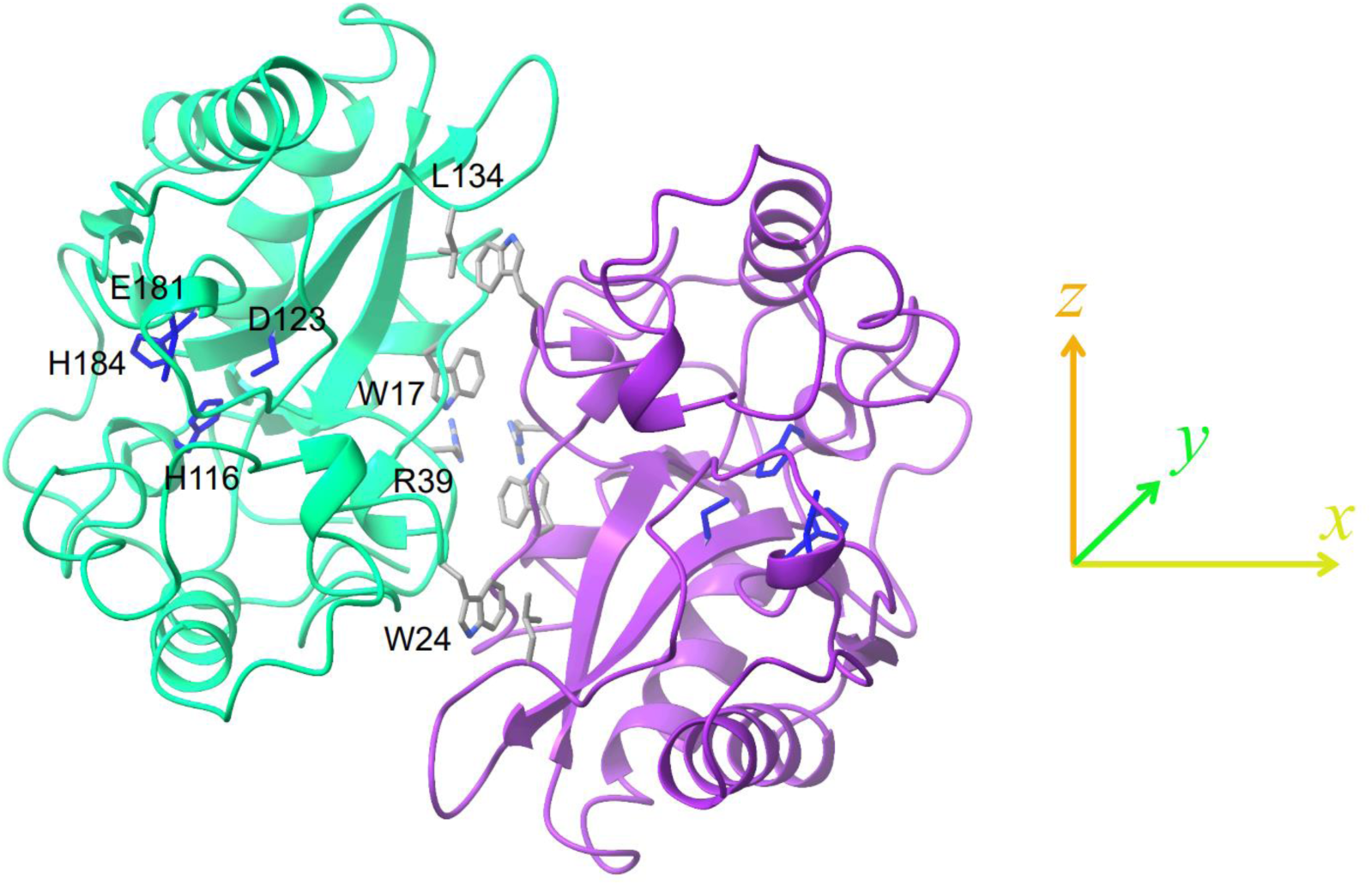
The dimeric structure calculated by Haddock (the same as in Figure S3). The main residues in the subunit-interaction region and the active site are shown. The coordinates on the right indicate the orientation of the alignment tensor.

**Figure S4.**
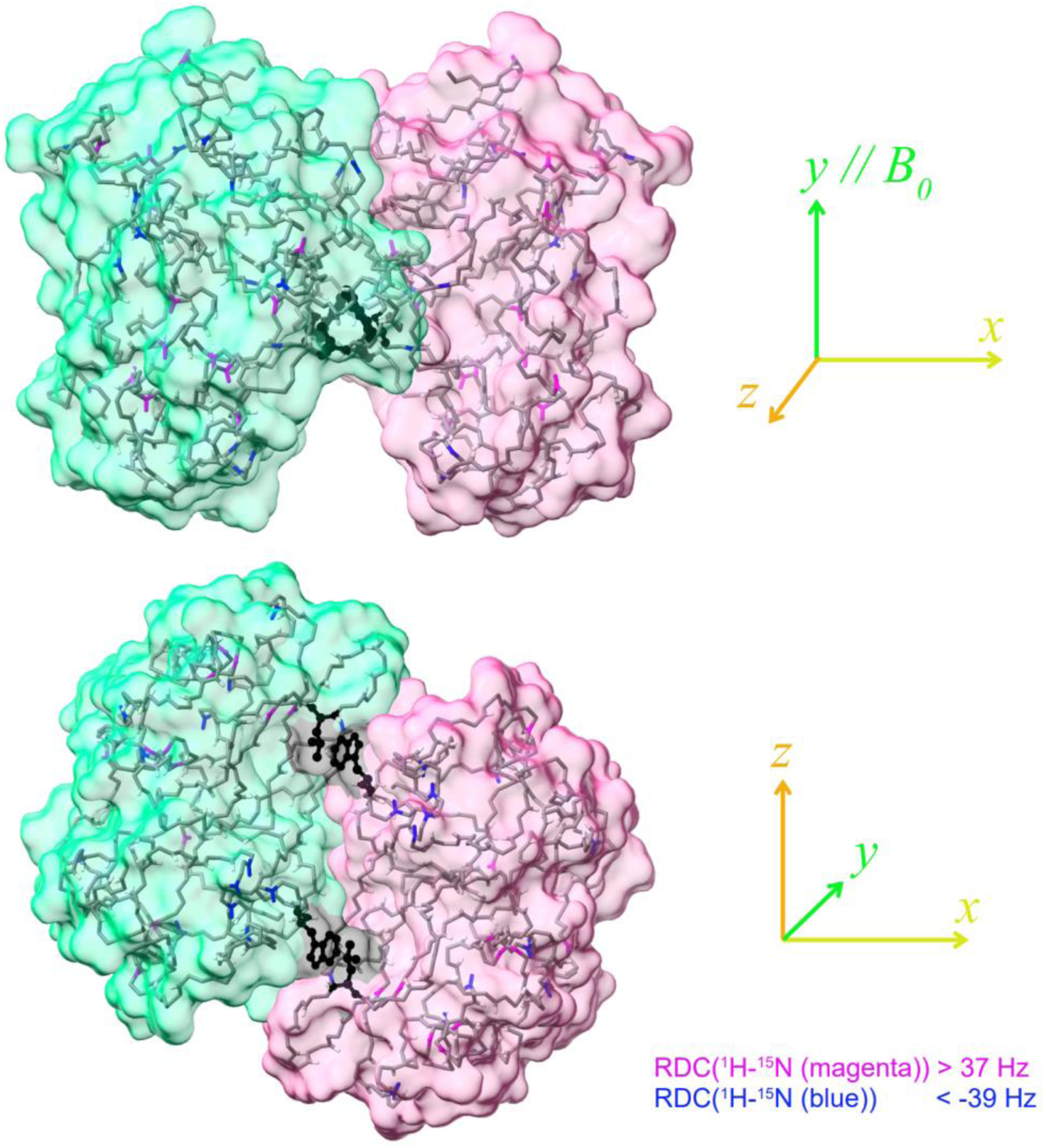
The dimeric structure calculated by Haddock based on RDC, TCS, and NOE data. Pales estimated the magnitude of the alignment tensor to be (*A*_xx_, *A*_yy_, *A*_zz_) = (1.01e-3, 1.53e-3, −2.54e-3). The tensor orientation provided by Haddock is shown on the right side of each structure diagram. For VanX, the *c*_2_ symmetry axis, *A*_yy_, was shown to be parallel to the static magnetic field. Amide bonds with RDC greater than 37 Hz are shown in magenta, and those with RDC smaller than −39 Hz are shown in blue. Side chains of Trp24 and Leu134, between which NOEs were observed, are shown in black.

